# Demographic change in an iconic dolphin population driven by habitat degradation following a marine heatwave

**DOI:** 10.64898/2026.09.25.754421

**Authors:** Felix Smith, Dominik M. Behr, Manuela R. Bizzozzero, Simon J. Allen, Richard C. Connor, Stephanie L. King, Arpat Ozgul, Michael Krützen

## Abstract

Extreme climate events are rapidly reshaping ecosystems and contributing to biodiversity loss, yet their long-term impacts remain poorly understood, particularly in marine systems. In 2011, an unprecedented marine heatwave (MHW) on the west coast of Australia drove declines in habitat-forming plants and across taxa spanning multiple trophic levels. Shark Bay, home to an iconic Indo-Pacific bottlenose dolphin (*Tursiops aduncus*) population, endured temperatures 3-5°C above long-term means, triggering catastrophic seagrass loss.

Using 20 years (2003-2023) of dolphin photo-identification data, we fitted a Bayesian Hidden Markov Model to estimate annual, age-class specific survival probabilities, abundance and recruitment rate, and their association with MHW-mediated seagrass loss in the western (WSB) and eastern (ESB) gulfs of Shark Bay.

Adult survival declined markedly, from 0.99 (0.98-1.00) pre-MHW to 0.85 (0.79-0.90) post-MHW in WSB, and from 0.96 (0.92-0.98) to 0.89 (0.85-0.95) in ESB, remaining below pre-MHW levels for several years, while juvenile and calf survival declined in both gulfs to a lesser extent. Recruitment declined in both gulfs and, although recovery was evident in WSB from 2019 onward, continued to decline in ESB until the end of the study. These sustained demographic declines drove substantial reductions in population growth and abundance in both WSB (39%) and ESB (32%). Demographic changes were associated with seagrass loss, here a proxy for habitat and prey availability, with clearest support for adult survival in WSB and a two-year-lagged recruitment response in ESB.

This pathway is likely to operate across diverse ecosystems as climate change drives increasing loss of habitat-forming species. The severity and duration of these demographic impacts illustrate that habitat change can cascade through trophic systems to overwhelm demographic resilience in upper-trophic species. This has negative implications for even long-lived, K-selected species, whose slow demographic recovery may not keep pace with the accelerating frequency and intensity of climate disturbance.

## 1. Introduction

Extreme climatic events are increasingly recognized as key drivers of ecological change and may have greater environmental impacts than slow changes in mean climate (Bailey and Van De Pol 2016; Gutschick and BassiriRad 2003; Harris et al. 2018; Jentsch et al. 2007; Maron et al. 2015; Maxwell et al. 2019; Smith 2011). For example, riverine forests in south-east Australia exhibited declining health in response to reduced river flows and falling ground water levels driven by water extraction, declining rainfall and increasing temperature (Mac Nally et al. 2011). Yet the most severe impacts occurred during acute drought and heatwave events, which caused dieback across 70% of the forest (Horner et al. 2009). Such examples illustrate that, while gradual environmental change may predispose ecosystems to stress, extreme events can drive the most severe and rapid ecological impacts. Extreme events are often short in duration but can expose organisms and populations to environmental conditions that exceed their capacity to acclimate to a greater extent than gradual changes associated with long-term climate trends (Gutschick and BassiriRad 2003).

Extreme weather and climate events, including storms (Lindroth et al. 2009), tropical cyclones (Feehan et al. 2024), droughts (Lewis et al. 2011), floods (Lake 2000), cold snaps (Garrett et al. 2022), fires (Legge et al. 2022) and heatwaves (Baum et al. 2026), can drive steep population declines, destabilize population dynamics, rapidly alter ecosystems and reshape patterns of biodiversity (Berger et al. 2018; Coulson et al. 2001). These events have caused mass mortalities (Breshears et al. 2005), reduced fitness and altered behavioural patterns through direct physical forcing (Matich and Heithaus 2012), as well as indirectly impacting species through rapid changes to habitats and resource availability (Nowicki et al., 2019; Wernberg et al., 2013).

Marine ecosystems are undergoing rapid change, with upper-ocean temperatures warming significantly across most regions during the last century (Calvin et al. 2023; Giralt Paradell et al. 2025). This surface warming is amplified by increasing upper-ocean stratification, which inhibits heat diffusion to depth (Capotondi et al. 2024). Against this backdrop, extreme climate events, such as marine heatwaves, are becoming more frequent, more intense and longer lasting (Oliver et al. 2018), with record breaking marine heatwaves documented in most ocean basins over the past decades (e.g., Benthuysen et al., 2018; Bond et al., 2015; Chen et al., 2014; Di Lorenzo and Mantua, 2016; Pearce et al., 2011; Smale et al., 2019; Sparnocchia et al., 2006). Marine heatwaves are driven by a combination of local oceanic heat fluxes and atmospheric conditions (Oliver et al. 2021), which are increasingly modulated by anthropogenic warming (Capotondi et al. 2024). Long-term increases in mean ocean temperatures have contributed substantially to observed increases in marine heatwave frequency and duration (Oliver et al. 2018), with 87% of marine heatwaves today attributable to human-induced warming (Frölicher et al. 2018). Given the projections of upper-ocean warming, marine heatwaves are expected to increase in magnitude and frequency throughout the twenty-first century (Meehl and Tebaldi 2004).

Marine heatwaves have had widespread and often calamitous impacts on marine ecosystems (Collins et al. 2019). Acute temperature changes have driven substantial ecological impacts, including the sustained loss of seagrasses (Wernberg et al. 2016, 2013), coral bleaching (Hughes et al. 2017), mass mortality across trophic levels (Gálvez et al. 2023; Oliver et al. 2017; Pearce et al. 2011), and large-scale range shifts of numerous species (Cavole et al. 2016; Oliver et al. 2018), altering the structure and function of entire ecosystems. Habitat compression and shifts in prey distribution may reduce prey availability and, in turn, individual fitness of upper-trophic species (Gálvez et al. 2023; Santora et al. 2020). Consistent with these mechanisms, marine heatwaves have been associated with population declines in humpback whales (Gabriele et al. 2022), Guadelupe fur seals (Gálvez et al. 2023) and other marine mammals (e.g., Nowicki et al., 2019).

Quantifying the impacts of marine heatwaves over ecologically meaningful timescales remains challenging because many studies lack long-term datasets and the unplanned nature of events means there is often limited data on the study system pre-event (Bailey and Van De Pol 2016). Thus, few studies have investigated the long-term response and lasting consequences of extreme events (Bailey and Van De Pol 2016). Addressing this gap is critical for predicting how increasingly frequent and intense marine heatwaves will shape marine megafauna populations in the coming decades.

In the Austral summer of early 2011, an unprecedented marine heatwave impacted the western coast of Australia (Feng et al. 2013). This event was driven by extraordinary La Niña conditions that increased the flow of the Leeuwin Current and transfer of warm tropical water southward (Feng et al. 2013). The Shark Bay World Heritage Site, a shallow embayment comprised of western and eastern gulfs bisected by the Peron Peninsula, was subject to extreme seawater temperatures for over 10 weeks. Temperatures recorded 3°C above long-term monthly means, reaching 30°C in some locations (Pearce and Feng 2013; Strydom et al. 2020; Thomson et al. 2015). The heatwave devastated habitat-forming seagrass meadows (Pearce et al. 2011), with a total of 1,310 km^2^ (∼31 %) lost, the single largest loss of seagrass biomass globally (Strydom et al. 2020; Thomson et al. 2015; Wernberg et al. 2013).

Seagrasses are marine foundation species that strongly influence ecosystem structure and function, providing key ecosystem services such as habitat provision, biodiversity support, nutrient cycling, sediment stabilisation and carbon sequestration (Barbier et al. 2011; Duarte 2002; Nordlund et al. 2016; Waycott et al. 2009). Pre-heatwave seagrass assemblages in Shark Bay were dominated by temperate species, with *Amphibolis antarctica* accounting for approximately 85% of seagrass cover (Fraser et al. 2014). The structural complexity of *A. antarctica* supported enhanced epifauna and epiphyte growth (Edgar and Robertson 1992), high invertebrate abundance (Wells et al. 1985), and provided refuge from predators (Hyndes et al. 2003), creating critical habitat for many fish species.

The loss of seagrass following the heatwave, including a 43% decline in *A. antarctica* cover (Strydom et al. 2020), led to reduced size and increased fragmentation of seagrass patches, substantially lowering habitat quality for dependent fauna (Strydom et al. 2026). Seagrass loss was associated with declines in abundance of seagrass-dependent megafauna throughout Shark Bay (Nowicki et al. 2019), compounding direct heatwave-driven mass mortality of invertebrate and fish communities across the region (Pearce et al. 2011). A ∼40% decline in benthic fish biomass in shallow habitats and ∼27% in deeper habitats accompanied the seagrass loss (Nowicki et al. 2019), substantially reducing prey availability for upper trophic predators. Consistent with this, survival and reproductive rate of Indo-Pacific bottlenose dolphins (*Tursiops aduncus*) in the western gulf of Shark Bay declined significantly following the heatwave (Wild et al. 2019). Given their slow development, late maturation and low reproductive rates (Mann et al. 2000), bottlenose dolphins are at particular risk from the prolonged effects of extreme climate events (Maxwell et al. 2019). However, the long-term demographic consequences and extent of population-level impacts across both gulfs remain unquantified.

Shark Bay provides an unparalleled opportunity to investigate the long-term impacts of extreme climatic events on the demographic parameters of a long-lived, slow reproducing and behaviourally flexible vertebrate. Seagrass loss was heterogenous between the gulfs (Strydom et al. 2020), the movement of dolphins between which appears negligible (Krützen et al. 2004). The two gulfs can therefore be treated as independent demographic sub-populations, providing a natural comparison of demographic responses under differing habitat conditions. Shark Bay Dolphin Research has studied the population since 1982, providing a multi-decadal dataset that spans both pre- and post-heatwave periods.

Using twenty years of photographic mark-recapture data, we applied a Bayesian Hidden Markov Model to quantify the long-term demographic consequences of the 2011 marine heatwave on Shark Bay’s dolphins. Specifically, we estimated changes in annual, age-class-specific survival and recruitment in both gulfs, examining the influence of sex, foraging strategy and seagrass dynamics. In doing so, we provide robust demographic parameter estimates to inform future assessments of the long-term viability of this population under increasingly frequent and intense marine heatwave disturbance.

## 2. Methods

### 2.1. Study Area

Shark Bay, a UNESCO-listed world heritage site, is located approximately 850km north of Perth on the west coast of Australia. It is a large (15,000km^2^) semi-enclosed bay divided into two gulfs by the Peron Peninsula (Fig. 1A). Both gulfs are shallow, with a mean depth of 9m, and characterized by extensive seagrass beds and sandflats divided by deeper channels (Sargeant et al. 2007; Tyne et al. 2012). There is limited movement of individuals between the two study sites, separated by ∼110km by water, indicating they function as separate sub-populations (Krützen et al. 2004; Marfurt et al. 2024).

**Figure 1:**
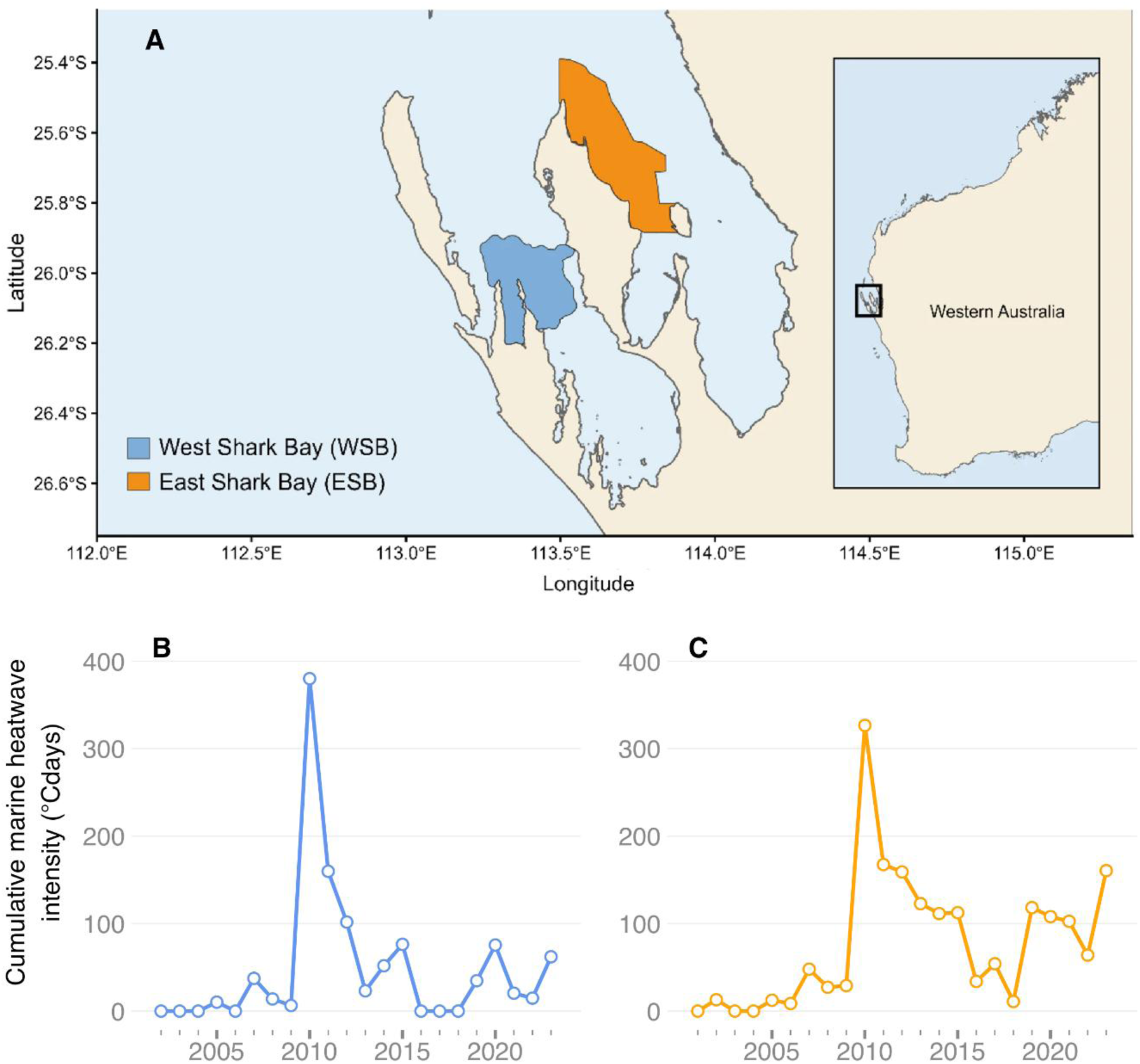
Study area and temporal variation in cumulative marine heatwave intensity in Shark Bay, Western Australia. A: Map of study sites in West Shark Bay (WSB) and East Shark Bay (ESB). Illustrated study site boundaries are based on a combination of habitat type (Sutton and Shaw 2020; modified in Bizzozzero et al. 2025), benthic features and the 95% kernel density estimate of all surveys included in the analysis; B and C: Annual cumulative marine heatwave intensity (°Cdays) in WSB and ESB, respectively. Cumulative marine heatwave intensity represents the sum of daily temperature anomalies (°C) above a fixed 30-year climatological threshold across all days classified as part of a marine heatwave event within each year (see Supplementary Methods S1 for estimation methods).

**Figure 2:**
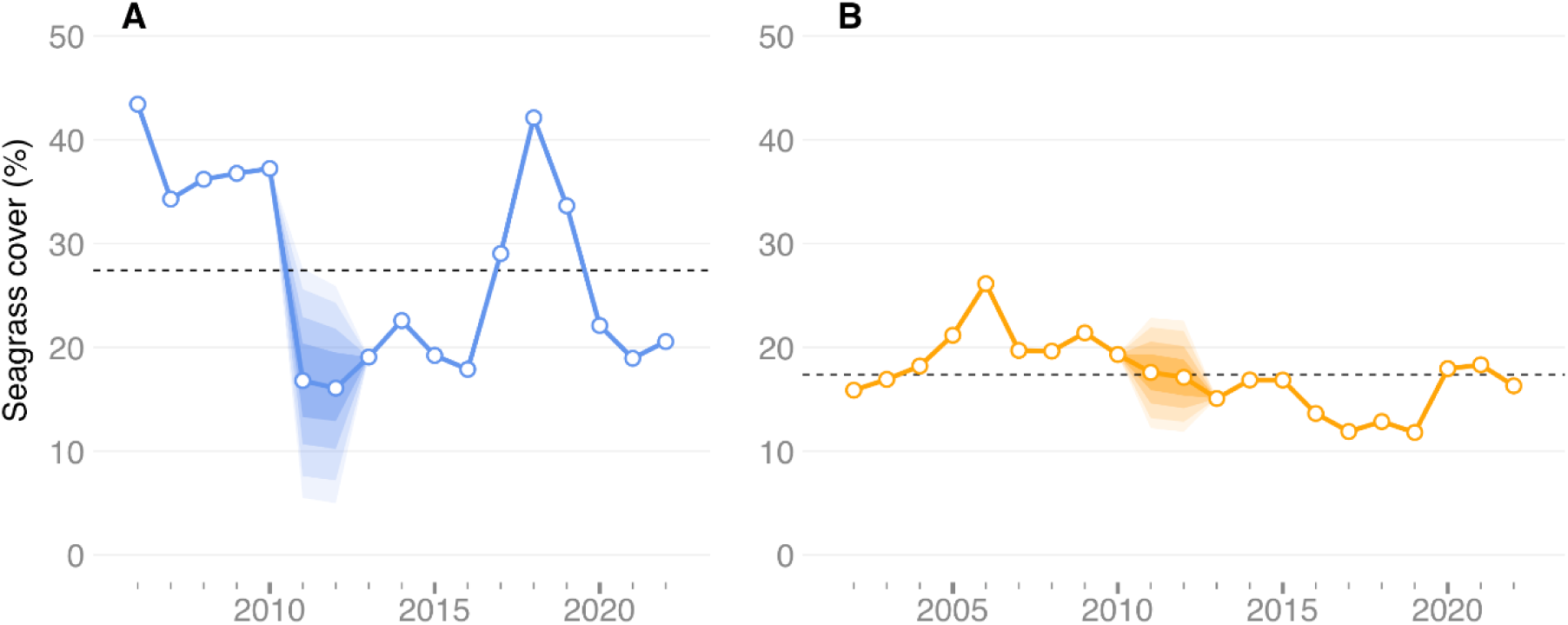
Annual seagrass cover (%) in (A) West Shark Bay (WSB) and (B) East Shark Bay (ESB). Imputed values from the seagrass sub-model are within graduated posterior credible interval ribbons (50%, 75%, 90%, 95%). Horizontal dashed line shows overall mean cover in each gulf.

### 2.2. Data Collection

We conducted boat-based surveys between June and October from 2007 to 2023 in western Shark Bay (WSB) and from 2003 to 2023 in eastern Shark Bay (ESB). Weather-dependent sampling (daylight hours, low wind, no rain, Beaufort Sea scale ≤ 3) either followed pre-determined, systematic transects or was conducted opportunistically within the confines of the study area (Nicholson et al. 2012; Connor and Krützen 2015).

When dolphins were sighted, we approached to conduct a photo-identification and behavioural survey, recording GPS location, water depth and temperature (O’Brien et al. 2020). Individual foraging strategy (‘sponger’ or ‘non-sponger’; Smolker et al., 1997) was also recorded. Sponging is a culturally transmitted foraging behaviour in which dolphins use marine sponges as a foraging tool (Krützen et al. 2014). Previous analysis suggested this foraging strategy, or at least the niche the strategy affords access to, provided some buffering from the demographic consequences of the heatwave (Wild et al. 2019), motivating its inclusion as a covariate in this study. This covariate therefore also serves to distinguish between competing pathways: a buffering effect may indicate prey declines specific to seagrass-associated habitat, whereas its absence may point to a broader, cross-habitat decline in prey availability. A dolphin was classified as a sponger after being observed foraging with a sponge on at least two separate occasions on different days (Kopps et al. 2014). Foraging strategy was confirmed for all individuals, with 7% (n = 54) and 3% (n = 39) of WSB and ESB study populations classified as spongers, respectively.

#### 2.2.1. Photo-identification of bottlenose dolphins

By comparing field photographs with our long-term catalogue, we identified individual dolphins, following well established protocols (Würsig and Würsig 1977). We identified individuals using distinctive dorsal fin features such as nicks, notches, fin shape and scarring patterns (Wursig and Jefferson 1990). Given our extensive archive for these sub-populations, we also made matches using more subtle or temporary markings. To ensure correct identification of individuals, a second observer independently verified all photo-identification data. In total, 780 and 1,193 individual dolphins were identified across the study period in WSB and ESB, respectively.

#### 2.2.2. Date of birth estimation

We estimated date of birth of dependent calves based on body size, presence of foetal lines and time since the last encounter of the mother in the absence of a calf (Mann and Smuts 1999). For individuals that were first seen as juveniles or adults, we based age estimates on body size, speckling patterns and date of first calf or onset of consorting behaviour (Connor and Krützen 2015; Krzyszczyk and Mann 2012; Smolker et al. 1992). For each individual, we assigned estimated birthdates with associated uncertainty intervals (in days) to reflect the accuracy of the estimate. Here, we defined age-classes as: calf (<3 years), juvenile (3–12 years), or adult (>12 years). While individuals in Shark Bay show variation in life-history timing (e.g., weaning age, age at first reproduction; Mann et al. 2000; Karniski et al. 2018), these transitions are not observed for all individuals. We therefore used fixed age thresholds to ensure consistent classification across the dataset, providing the discrete age-classes required by the HMM.

#### 2.2.3. Sex estimation

We identified sex either via direct observations of the genitals (Smolker et al. 1992), genetically (based on Gilson et al., 1998, modified in Bizzozzero et al., 2019; Marfurt et al., 2022), through the presence of a dependent calf (Connor and Smolker 1985) or behaviourally by repeated observations of male alliance-typical behaviour, i.e., regularly travelling side-by-side, engaging in synchronous surfacings, consorting of females and intergroup aggression with other males (Connor and Krützen 2015). For 50% of individuals in WSB (n = 388; 200 females, 188 males) and 72% of individuals in ESB (n = 859; 417 females, 442 males), we confirmed sex, with the remainder unsexed (n = 392 and n = 334, respectively).

### 2.3. Seagrass Coverage as an Environmental Covariate

To examine the potential environmental drivers of demographic variation, we included annual seagrass coverage (km^2^) as a covariate, serving as an index of the magnitude and persistence of the heatwave’s ecological impact, and thus of prey availability. Dolphins in Shark Bay forage across a mosaic of habitat types, including seagrass meadows, sand flats, and deep channels, that support different prey communities (Bizzozzero et al. 2026). Pre-heatwave, seagrass meadows characterised much of the shallow habitat in Shark Bay, including areas of high prey biomass (Heithaus and Dill 2002), and supported dolphin foraging on seagrass-associated fish species (Nowicki et al. 2019). Following the heatwave, the composition of this mosaic shifted, with reduced seagrass cover accompanied by increases in bare sand and greater habitat fragmentation (Strydom et al. 2026). Numerous fish species occupying reef and rocky shoreline habitats also use seagrass as nursery areas (Guidetti 2000), meaning seagrass-independent foraging strategies may be indirectly affected through prey dependence on seagrass nursery habitat. The heatwave is also likely to have affected fish directly through thermal stress (Pearce et al. 2011), thus we do not consider seagrass coverage as the sole pathway linking the heatwave to prey decline. Rather, it provides a biologically meaningful, quantitative index of the event’s ecological impact that captures the prolonged degraded state and slow recovery of the ecosystem following the heatwave.

#### 2.3.1. Estimation of annual seagrass coverage

We derived annual seagrass coverage (km²) from Landsat satellite imagery (2001–2023, excluding 2011–2012 – *see below*), using a random forest classifier applied to annual composite images in Google Earth Engine following methods detailed in Bizzozzero et al., (2025). Full details of image selection, classification, and validation procedures are provided in Supplementary Methods S2.

Annual seagrass coverage was not available for 2011 and 2012 in either gulf due to data availability constraints caused by reduced satellite image quality during this period. As marine heatwaves are a known driver of seagrass loss in Shark Bay (Strydom et al. 2020), we used a cumulative marine heatwave intensity covariate (cMHW; °Cdays; Hobday et al. 2016; See Supplementary Methods S1) to infer seagrass coverage for years with missing data via an autoregressive sub-model described in Supplementary Methods S2.2. The sub-model was embedded within the demographic model, allowing uncertainty in imputed seagrass values to propagate through to the vital rate estimates.

### 2.4. Data Analysis

#### 2.4.1. Robust design sampling strategy

We estimated age-class specific detection and demographic parameters (annual apparent survival probabilities, annual recruitment, temporary emigration, annual abundance and population growth rate) and their association with sex, foraging strategy and seagrass dynamics using a Bayesian Hidden Markov model (HMM) implemented with a closed robust design sampling strategy (Bond et al. 2025; Gibson et al. 2018; Rankin et al. 2016; Riecke et al. 2018). Survival probability represented the probability of an individual alive in year *t* surviving to year *t* + 1; recruitment rate represented the number of calves entering the population in year *t* relative to the number of adult females alive at year *t* - 1, where calves have survived long enough to be observed in the study; and abundance was defined as the total number of dolphins alive at time *t*.

In the robust design, primary periods (here, field seasons) comprise >1 nested capture occasions (secondary periods). Primary periods occurred once per year (usually June–October) and were separated by at least seven months. Each primary period was divided into multiple two-week secondary periods. This was selected as the best compromise between model performance and the assumption that secondary sampling occasions should be instantaneous (Williams et al. 2002; see Supplementary Methods S3 for consideration of other assumptions). During each primary period, the population is assumed to be demographically closed: individuals remain “onsite” (within study site and available for detection) or “offsite” (temporarily outside the study site or within study site but unavailable for detection), with no births or deaths occurring. This structure increases effective detection probabilities for animals that are difficult to observe and supports the estimation of temporary emigration (Kendall and Nichols 1995). Between primary periods, the population is assumed to be “open”, allowing individuals to transition between “onsite” and “offsite” states, and births and deaths to occur. We constructed capture histories for each individual, recording whether the animal was detected (1) or not detected (0) during each secondary period within a primary sampling period. The HMM models individual state dynamics as latent variables, allowing transitions between biological states to be inferred even when individuals are not observed. See Supplementary Methods S4 for details.

#### 2.4.2. Accounting for unequal survey effort

To account for unequal spatial survey effort, we subdivided each study area into survey zones based on a combination of habitat type (Bizzozzero et al. 2025) and focal areas typically targeted on a given survey day (seven in ESB, eight in WSB). We used the proportion of days each zone was surveyed within each secondary period as a continuous effort covariate on detection probability. Full details are provided in Supplementary Methods S5.

#### 2.4.3. Assessment of Hidden Markov Model Goodness of Fit

To assess goodness of fit of our Bayesian HMM, we performed posterior predictive checks by generating replicated datasets from the posterior predicted distribution and comparing them to the observed data (Gelman et al. 2013). We compared the number of individuals detected during each primary period in the replicated data against the observed counts and derived Bayesian *p-*values. These represent the probability that the replicated data are more extreme than the observed data. Values near 0.5 represent a good model fit (Gelman et al. 2013) and those outside the 0.10–0.90 range indicate poor fit (Hobbs and Hooten 2015).

#### 2.4.5. Identification of Demographic Processes Influencing Population Dynamics

We conducted a retrospective population analysis to identify which demographic processes most strongly influenced population dynamics (Schaub et al. 2013; Schaub and Kéry 2021). To do so, we quantified the contribution of the annual variation of each parameter by calculating its correlation with annual population growth rate. We calculated Pearson correlation coefficients for each posterior sample and reported mean values, along with 95% credible intervals, using R version 4.4.2 (R Core Team 2024; Schaub et al. 2013).

### 2.5. Hidden Markov Model Fitting and Inference

We coded and fitted the Bayesian HMM in NIMBLE using version 1.3.0 of the R package *nimble* (De Valpine et al. 2024). We used non-informative priors for all parameters. To perform the Markov chain Monte Carlo (MCMC) sampling, we used six parallel chains of 150,000 iterations each, with a burn in phase of 50,000 iterations and every 10th sample retained for thinning. We assessed convergence by visual inspection of trace plots and by computing the Gelman-Rubin convergence statistic, *Ȓ* (Gelman and Rubin 1992), with values <1.1 indicating satisfactory convergence.

## 3. Results

Following the 2011 marine heatwave, both dolphin sub-populations suffered pronounced and prolonged declines in demographic rates. Adult survival decreased in both gulfs, remaining depressed for several years (Fig. 3A, B). Juvenile and calf survival also declined, but patterns were less pronounced and more variable across years and gulfs (Fig. 3C, D, E, F). Recruitment declined in both gulfs, but while there was recovery in recruitment in WSB, ESB showed no sign of recovery in the study period (Fig. 4). These changes were reflected in sustained reductions in population growth rates and overall abundance in both sub-populations (Fig. 6). Reductions in demographic rates coincided with declines in seagrass coverage following the heatwave and only partial recovery in subsequent years (Fig. 5). Although both sub-populations showed declining trends in demographic rates, the magnitude and timing of demographic responses differed between the gulfs.

**Figure 3:**
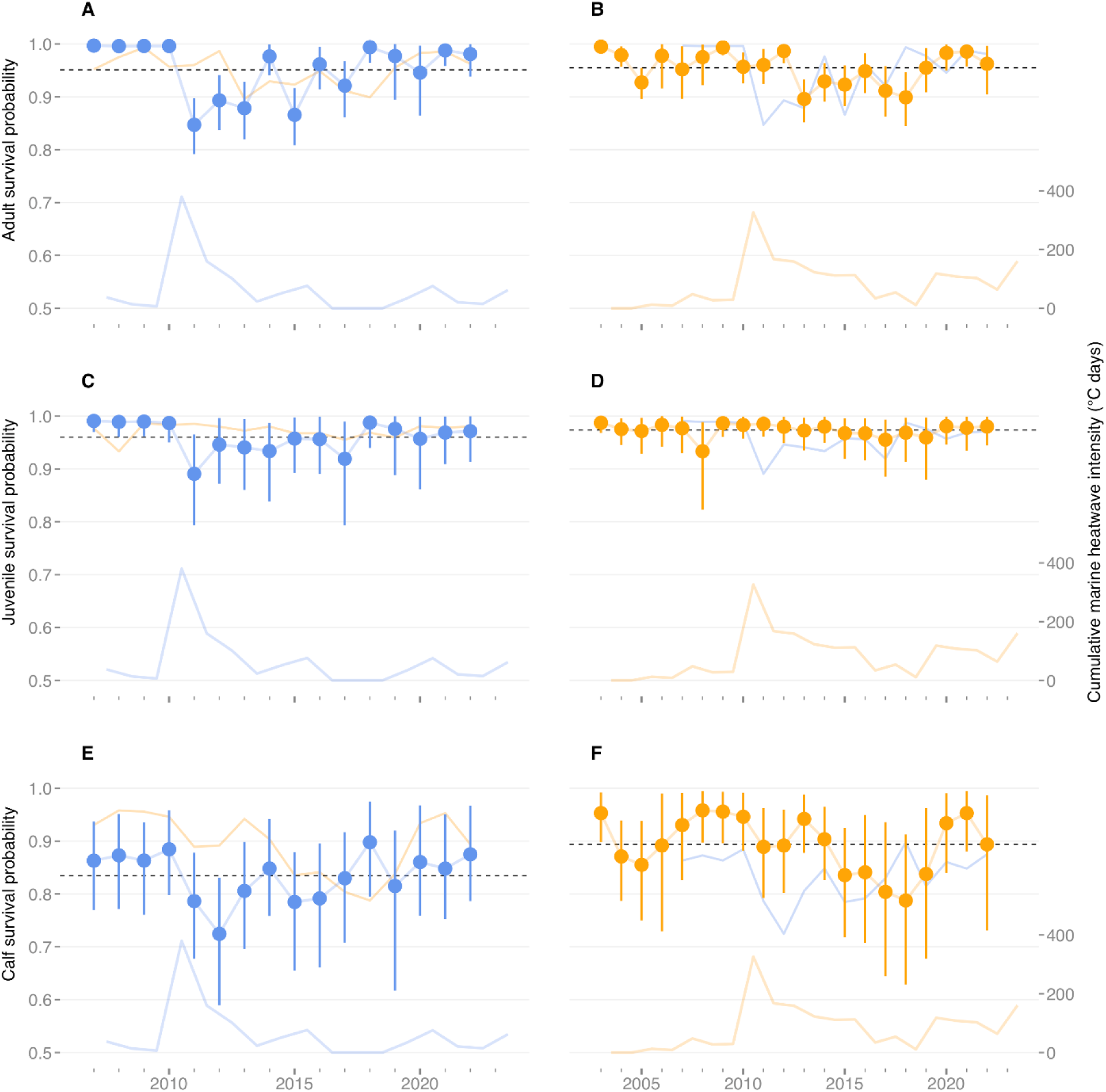
Annual age-class-survival probability for West Shark Bay (WSB; A, C, E) and East Shark Bay (ESB; B, D, F), showing adult (A, B), juvenile (C, D), and calf (E, F) survival probabilities. Points show posterior means; bars show 95% credible intervals. Horizontal dashed line shows overall mean survival probability for each age class and gulf. The background trend line in each panel shows the corresponding annual survival probabilities for the same age-class in the other gulf. The lower trendline in each panel shows cumulative marine heatwave intensity (°Cdays) for that gulf.

**Figure 4:**
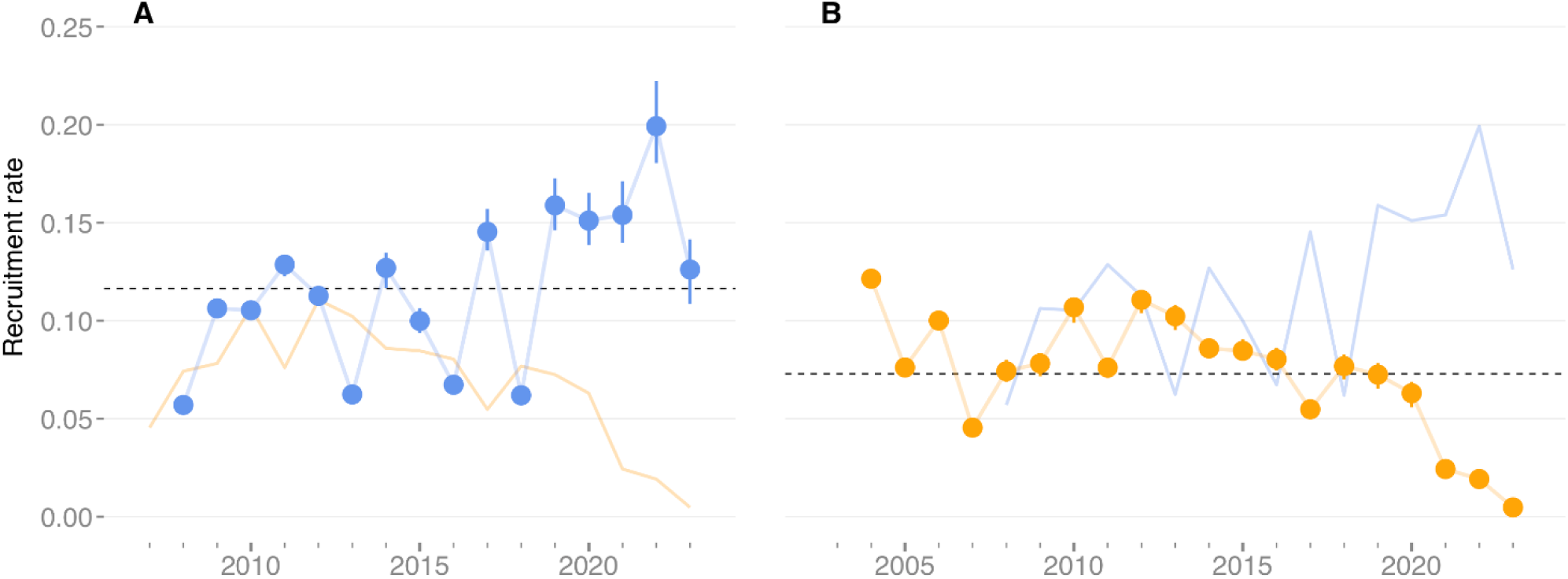
Annual recruitment rate for West Shark Bay (A) and East Shark Bay (B). Points show posterior means; bars show 95% credible intervals. Horizontal dashed line shows overall mean annual recruitment rate in each gulf. The background trend line shows annual recruitment rate for the other gulf.

**Figure 5:**
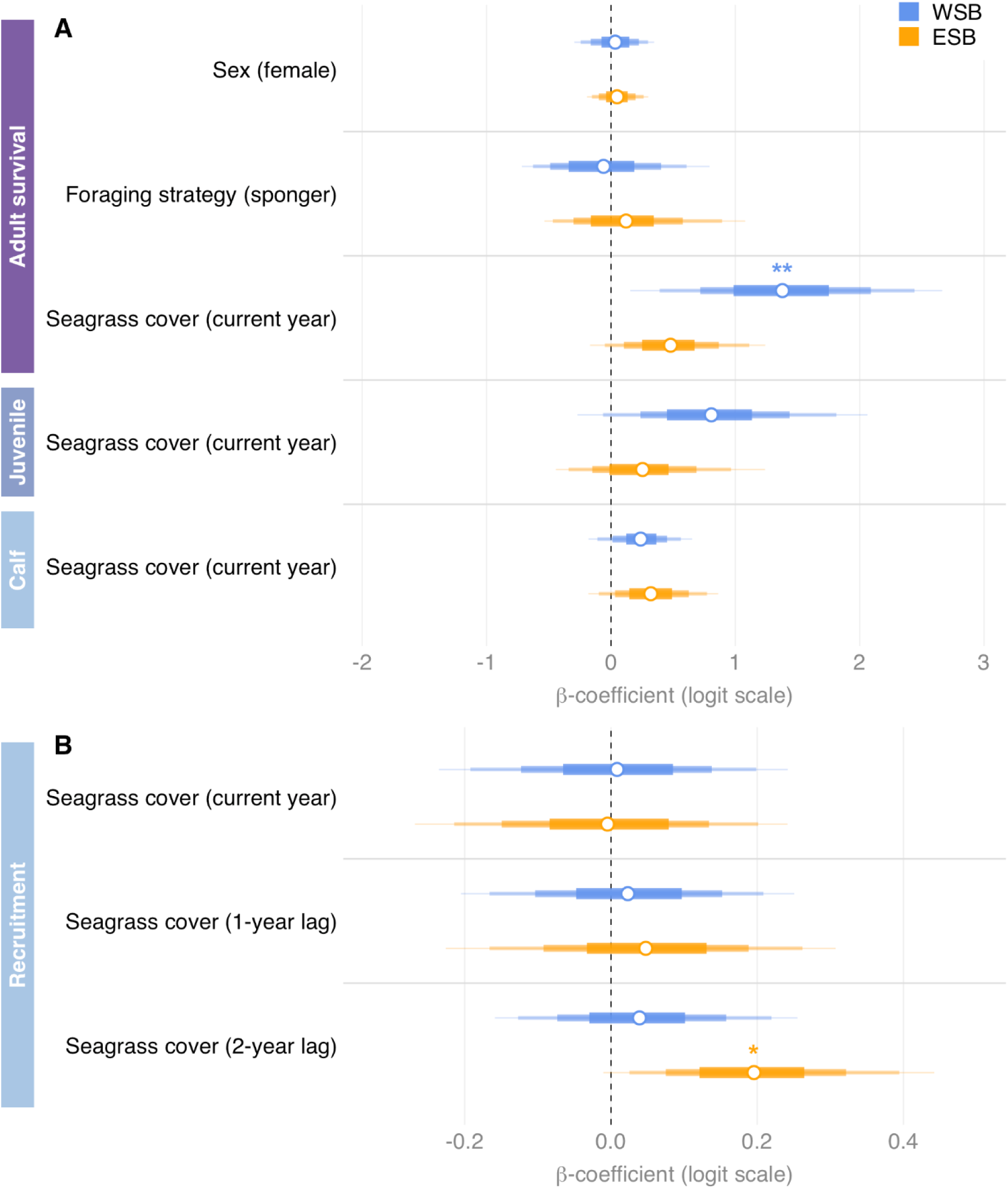
Forest plot of covariate effects (β-coefficient, logit scale) on survival probabilities and recruitment rate in West Shark Bay (WSB) and East Shark Bay (ESB). (A) Effects of sex, foraging strategy, and current-year seagrass cover on adult, juvenile and calf survival probability. (B) Effects of seagrass cover at three biological lags on recruitment rate. Seagrass cover (gestation; current year): food availability during the 12-month gestation period leading to the birth of calves counted in primary period *t;* Seagrass cover (conception; 1-year lag): food availability during the year of conception, reflecting conditions when females become pregnant; Seagrass cover (pre-conception; 2-year lag): food availability in the year prior to conception, reflecting maternal body condition before the reproductive attempt. Effects were estimated in separate models. Points show posterior means; bars show graduated credible intervals (50% thickest, 75%, 90%, 95% thinnest). Directional support is based on posterior probability of consistent sign: * = p > 0.90; ** = p > 0.95.

**Figure 6:**
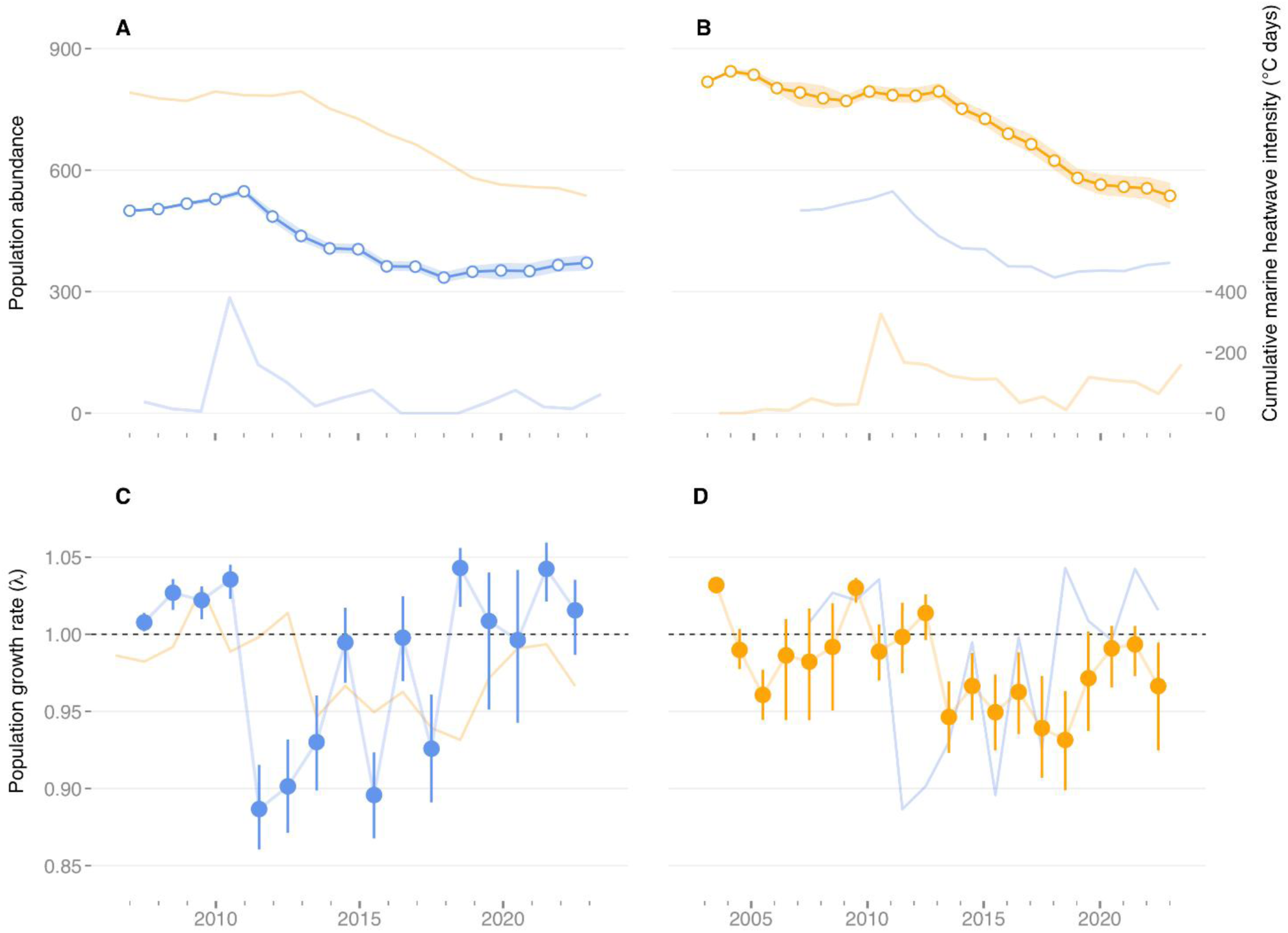
Total annual population abundance (onsite + offsite) in West Shark Bay (WSB; A) and East Shark Bay (ESB; B) Points show posterior means; ribbon shows 95% credible interval. Bottom trendline shows cumulative marine heatwave intensity (°Cdays). The background trend line shows dolphin abundance for the other gulf. Population growth rate (λ) in WSB (C) and ESB (D). Points show posterior means; bars show 95% credible intervals. Dashed black line at zero (log lambda = 0, lambda = 1) indicates a stable population; values above the dashed line indicate years of population increase and values below indicate years of decline. The background trend line shows annual population growth for the other gulf.

### 3.1 Estimates of Survival Probabilities

#### 3.1.2. Adult Survival Probabilities

Adult survival probabilities remained consistently high prior to the 2011 marine heatwave in WSB, but fell immediately after, to a minimum of 0.85 between 2011 and 2012 (95% CI: 0.79-0.90; Fig. 3A). Survival remained depressed for several years, showing some recovery between 2016 and 2017 but not consistently returning to pre-heatwave values until 2022. In contrast, adult survival in ESB did not decline until between 2013 and 2014 (0.89, 95% CI: 0.85-0.95), three years after the heatwave, and reduced estimates persisted until 2019 (Fig. 3B).

#### 3.1.3. Juvenile Survival Probabilities

Juvenile survival probabilities in WSB showed a similar trajectory, declining following the heatwave (2011-2012: 0.89, 95% CI: 0.80-0.96) and remaining depressed until 2019 (Fig. 3C). However, the decline in juvenile survival appeared to be smaller than that for adults. In ESB, juvenile survival probabilities remained relatively high, but a slight decline was observed, starting between 2015-2016 (0.97, 95% CI: 0.93-0.99). Survival probabilities remained below average until 2020 but, overall, were higher than for other age-classes (Fig. 3D).

#### 3.1.4. Calf Survival Probabilities

Calf survival in WSB declined immediately post-heatwave, reaching a minimum between 2012 and 2013 (0.73, 95% CI: 0.59-0.83; Fig. 3E). Survival remained below pre-heatwave levels for much of the following decade, with sustained recovery to consistently high levels only apparent from 2021 onwards, despite a brief increase between 2018 and 2019. Calf survival declined in ESB between 2015-2016, reaching a minimum between 2018-2019 (0.79, 95% CI: 0.63-0.92; Fig. 3F) and returning to pre-heatwave levels between 2020-2021.

### 3.2. Estimates of Annual Recruitment Rate

Annual recruitment patterns differed between gulfs: they remained consistent in WSB between 2009 and 2012 (range: 0.101-0.123), following a year of low recruitment in 2008. Recruitment declined markedly in 2013 (0.062), two years after the heatwave, and remained low until 2016, with posterior distributions centred below the long-term mean (Fig. 4A). Recruitment increased in 2017, but this was not sustained in 2018. A consistent recovery was observed only from 2019 onwards, with recruitment remaining elevated in subsequent years (range: 0.109-0.180). In ESB, recruitment was, overall, lower than in WSB and showed more temporal variation pre-heatwave (range: 0.045-0.121). The impact of the heatwave on recruitment appeared to be delayed, with a steady decline in recruitment from 2014 (0.081) until the end of the study (2023: 0.005; Figure 4B), reaching near-zero levels with no signs of recovery to pre-heatwave levels.

### 3.3. Association of Seagrass Coverage, Sex and Foraging Strategies with Survival and Recruitment Declines

Seagrass coverage in WSB had a credible positive effect on survival, with survival probability rising alongside seagrass coverage across the gulf (Fig. 5A). This was particularly true for adults (β = 1.35, 95% CI: 0.20, 2.81), while the effect was weaker and less certain for calves and juveniles. In ESB, the effect of seagrass coverage on adult survival was positive, but not credible (β = 0.49, 95% CI: -0.31, 1.50; Fig. 5A), with no clear effects for calves and juveniles. Sex and foraging strategy (sponger vs. non-sponger) had no detectable effect on adult survival in either gulf.

There was no detectable effect of seagrass coverage on per-capita recruitment in WSB for any of the three biological lags tested (Fig. 5B). The direction of association was consistently positive across biological lags, however, with posterior probabilities of a positive effect ranging from 54% (seagrass coverage at time *t*; gestation) to 65% (seagrass coverage at time *t*-2 years; pre-conception), indicating that increased seagrass coverage had a weak but consistently positive effect on recruitment. Seagrass coverage in ESB two years prior to the survey year (*t*-2; pre-conception) showed a clear positive effect on per-capita recruitment (β = 0.19, 95% CI: −0.02, 0.43, PP = 96.5%: Fig. 5B), while this was not the case for time *t* and *t*-1 (gestation and conception, respectively).

Overall, recruitment was primarily driven by unexplained year-to-year variation, likely reflecting unmeasured environmental drivers, stochastic variation in female reproductive success, or fluctuations in calf survival during the early dependency period not captured by the seagrass covariate alone. Across all lag models, in both gulfs, the random year effect accounted for most of the modelled year-to-year variation in recruitment. In WSB, seagrass coverage explained 11%, 10% and 14% of year-to-year recruitment variability at the current-year, one-year and two-year lags, respectively, with the remaining 86-90% captured by the random year effect. In ESB, seagrass coverage explained 7% and 8% at the current-year and one-year lags, respectively. The exception was the two-year lag model in ESB, where seagrass explained 28% of year-to-year recruitment variability, with 72% retained by the random year effect.

### 3.4. Estimates of Temporary Emigration Parameters

Temporary emigration in both WSB and ESB followed a Markovian process with strong state persistence (Table S4), indicating consistent patterns of site use, with individuals showing a high probability of remaining either available or unavailable for detection across years. In WSB, the probability of leaving the study area was low (*γ*′′ = 0.13), indicating strong site fidelity. However, individuals that were offsite had a high probability of remaining absent in subsequent primary periods (*γ*′ = 0.71), suggesting persistent temporary emigration. Similarly, in ESB there was a low probability of leaving the study area (*γ*′′ = 0.12) but high probability of remaining absent in subsequent primary periods if offsite (*γ*′ = 0.74). This pattern reflects predictable, repeatable space use, where some individuals are consistent residents of the study area while others are persistently peripheral, likely those whose home ranges only partially overlap the surveyed zone. As such, temporary emigration represents movement in and out of the boundaries of the study area, rather than wide ranging movement away.

### 3.5. Estimates of Abundance and Population Growth Rates

Dolphin abundance in WSB showed an upward trend between 2007 and 2011, with population growth rates λ ranging from 1.01-1.04 (Figs. 6A, C). Following the marine heatwave, growth rates were predominantly negative between 2012 and 2018 (λ = 0.89-0.93), resulting in a dramatic drop in population size from a pre-heatwave high of 548 dolphins (95% CI: 539-555) in 2011 to a low of 335 individuals (95% CI: 323-349) in 2018. From 2019 to 2023 growth rates were mostly positive (λ = 1.02-1.04), leading to a modest population recovery to 371 individuals (95% CI: 352-391) in 2023.

In ESB, dolphin abundance remained relatively stable between 2003 and 2013 (range: 771-844; λ ranging from 0.96-1.03), with no immediate change following the marine heatwave (Figs. 6B, D). However, from 2014 (752, 95% CI: 734-772) until 2023 (537, 95% CI: 504-569), the population showed a long, steady decline (λ range: 0.93-0.99) with no sign of recovery at the end of the study.

#### 3.5.3. Associations between vital rates and population growth

Population growth rates in WSB were strongly correlated with changes in adult survival (r = 0.93, 95% CI: 0.87, 0.97; Fig. 7). Calf survival, juvenile survival, and recruitment also showed positive correlations (r range: 0.56-0.60), although these relationships were weaker. Similarly, adult survival showed the strongest correlation with annual population growth in ESB (r = 0.88, 95% CI: 0.81, 0.93; Figure 7). Calf survival and recruitment (r range: 0.25-0.51) also showed positive correlations. Juvenile survival showed a weaker and more uncertain positive correlation (r = 0.32, 95% CI:−0.13, 0.66), with credible intervals broadly overlapping zero, suggesting limited detectable contribution to year-to-year variation in population growth rate in ESB. These results indicate that while multiple vital rates covary with population growth, adult survival was the vital rate most strongly associated with interannual variation in population growth in Shark Bay dolphins.

**Figure 7:**
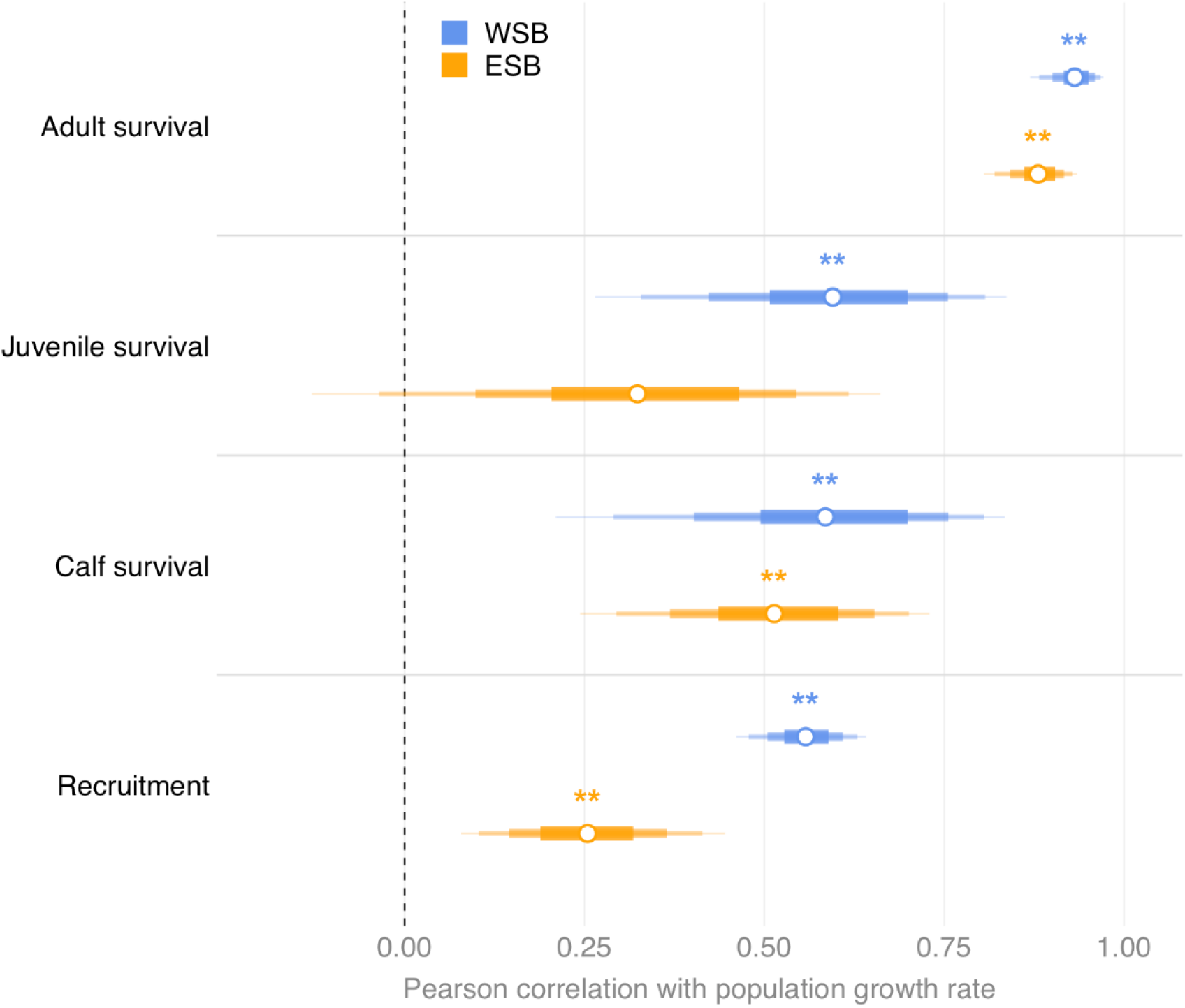
Posterior correlation between vital rates and annual population growth rate λ in West Shark Bay (WSB) and East Shark Bay ESB, computed for each MCMC sample across years. Points show posterior means; bars show graduated credible intervals (50% thickest, 75%, 90%, 95% thinnest). Directional support is based on posterior probability of consistent sign: * = p > 0.90; ** = p > 0.95. Year-specific survival estimates used in these correlations include the actual posterior seagrass values for each year; adult survival is computed for a reference individual (male, non-sponger), i.e. sex and foraging strategy effects are not applied.

### 3.6. Assessment of Model Fit

Posterior predictive checks suggested that the model broadly reproduced annual detection counts, with an average Bayesian *p*-value of 0.77 in WSB and 0.65 in ESB across all primary survey events (Supplementary Fig. S6). Although these averages lie above the theoretical optimum of 0.5, they were driven by a small number of boundary years rather than by systematic model misfit: the earliest survey years, in which individuals first identified later in the study are inferred to have been present earlier (an inherent feature of open-population models (Schwarz and Arnason 1996), and years with no survey effort (2007-2008 in ESB) or near-zero effort due to COVID-19 lockdowns (2020 in WSB; Supplementary Information S6).

## 4. Discussion

Sustained reductions in survival and recruitment rate following the 2011 marine heatwave drove sub-population declines of more than one-third in both gulfs of Shark Bay, demonstrating severe and long-lasting demographic consequences. There were, however, differential effects between gulfs – West Shark Bay (WSB) suffered an immediate and long-lasting decline in abundance before partial recovery, while the decline in East Shark Bay (ESB) was delayed but sustained through to the end of the study period. Declines in survival across all age-classes and in recruitment persisted for multiple years in both gulfs, although the observed impacts were delayed in ESB. These declines in survival and recruitment were closely tied to the heatwave’s impact on seagrass in Shark Bay. Recruitment recovered from 2019 onwards in WSB but continued to fall throughout the study period in ESB. The concurrent declines in survival and recruitment indicate positive covariance among vital rates, which amplifies population variability (Compagnoni et al. 2016; Morris et al. 2008) and likely drove the sustained abundance declines observed over the decade following the heatwave.

Adult survival probability declined in both gulfs following the heatwave, dropping from 0.99 to 0.85 in WSB in 2012 and from 0.96 to 0.89 in ESB in 2013. These reductions are comparable in magnitude to post-heatwave declines in humpback whales (*Megaptera novaeangliae;* Gabriele et al., 2022) and Steller sealions (*Eumetopias jubatus;* Hastings et al., 2023), as well as polar bears (*Ursus maritimus*) during extended ice free periods (Regehr et al. 2010). Although apparent survival estimates cannot distinguish mortality from permanent emigration (Lebreton et al. 1992), direct mortality associated with extreme heat events has been documented in a range of mammalian and avian taxa. Heatwave associated mortality has been recorded for Guadelupe fur seals (*Arctocephalus townsendi;* Gálvez et al. 2023), flying foxes (*Pteropus* spp.; Welbergen et al. 2008), howler monkeys (*Alouatta palliata mexicana*; Pozo-Montuy et al. 2024), koalas (Gordon et al. 1988) and humans (Mitchell et al. 2016), as well as the common murre (*Uria aalge*; Piatt et al. 2020), highlighting the capacity of extreme climatic events to drive substantial demographic impacts across diverse taxa. While abnormal dolphin mortality was not observed following the heatwave (Nowicki et al. 2019), the high philopatry, stable home ranges (Tsai and Mann 2013), habitat specialisation (Bizzozzero et al. 2026) and strong social bonds (Connor et al. 2022; Smolker et al. 1992) characteristic of Shark Bay dolphins suggest that permanent emigration in response to the heatwave is unlikely. This is further consistent with the marked genetic differentiation of Shark Bay dolphins from neighbouring populations to the north and south (Marfurt et al. 2024, 2026), reflecting limited historical gene flow and supporting the view that Shark Bay dolphins do not readily disperse to adjacent regions.

In slow growing, K-selected species, characterized by late maturation, small litter size and long life spans, population growth is often primarily influenced by adult survival (Crone 2001). This is evident in the Shark Bay population through the strong correlation between changes in adult survival and annual population dynamics. While populations may be resilient to short or intermittent periods of poor adult survival, sustained depression is likely to reduce population growth rate and drive severe population declines (Hastings et al. 2023).

We detected no effects of sex or foraging strategy on adult survival. Sex-differentiated survival in cetaceans is often attributed to the costs of male mating competition (Ralls et al. 1980). In Shark Bay, however, male alliance-based consortship competition (Connor and Krützen 2015) appears to reduce body condition similarly in both sexes (Cicciarella et al., 2026 *in prep*). Additionally, incomplete sex determination (50% unknown in WSB, 28% in ESB) likely limited our power to detect such an effect regardless. The lack of a foraging strategy effect contrasts with previous findings that sponging dolphins were somewhat buffered from the immediate effects of the heatwave (Wild et al. 2019). However, spongers comprised only a small fraction of each sub-population (7% in WSB, 3% in ESB), likely limiting both our statistical power to detect a strategy-specific survival effect and the ability of individual-level buffering to influence the population as a whole. Beyond this, sustained seagrass loss and the associated prey decline may have been too much for behavioural buffering to offset; alternatively this null result may also reflect a broader effect of the heatwave on prey across habitats and foraging niches. This effect may have been compounded by the high degree of individual foraging specialisations characteristic of Shark Bay dolphins, where individuals show surprisingly strong fidelity to their specific foraging niche (Bizzozzero et al. 2026). Species with such specialised habitat requirements are considered particularly vulnerable to extreme climate change (Maxwell et al. 2019), and individual foraging specialisations may therefore represent a constraint on behavioural buffering capacity when habitat loss is sustained and widespread.

Despite the expectation that adult survival is buffered relative to other vital rates in long-lived, K-selected species (Pfister 1998; Gaillard et al. 2000), adults showed large declines in both gulfs. Juvenile survival also declined in both gulfs, though less severely, an unexpected pattern, but one also observed in Steller sea lions following the north-east Pacific heatwave (Hastings et al. 2023). The greater vulnerability of adults may reflect the cumulative energetic costs of somatic maintenance and reproduction (McEntee et al. 2023), which may render adults particularly sensitive to prolonged energetic stress. The comparatively smaller decline in juvenile survival may be attributable to the absence of reproductive costs during the juvenile life stage or partial buffering through relaxed density-dependent competition following declines in adult abundance (e.g., López-Roig and Serra-Cobo 2014), although these mechanisms remain speculative.

Calf survival also declined in both gulfs following the heatwave. Calves, with underdeveloped dive physiology and limited foraging experience (Noren et al. 2002; Noren 2020), are dependent on their mothers, typically nursing until the ages of three to four years (Mann et al. 2000; Karniski et al. 2018). As such, calf survival reflects not only the calf’s physiological vulnerability but also the capacity of mothers to maintain reproductive investment under environmental stress. Declines in calf survival following extreme climate events have been documented in other cetacean species; humpback whale calf survival fell dramatically following the NE Pacific marine heatwave (Gabriele et al. 2022; Barber-Meyer et al. 2026), surpassing the impact documented in this study for both gulfs of Shark Bay, suggesting that calf survival may be broadly sensitive to acute environmental perturbation across cetacean species. The close coupling of calf survival and maternal condition is further reflected in recruitment patterns.

In both gulfs, recruitment declined post-heatwave and remained depressed for several years. While there was recovery in the western gulf, recruitment continued to decline to the end of the study in the eastern gulf. Seagrass loss was accompanied by a marked decline in benthic fish biomass (Nowicki et al. 2019), and energetic limitation is likely the proximate cause of the observed declines. However, the specific physiological mechanisms cannot be determined from our data. Energetic limitation may increase rates of reproductive failure through abortion or neonate mortality if the nutritional needs of the mother and offspring cannot be met (Trites and Donnelly, 2003). Delays in reproductive maturity and suppressed ovulation may also occur if food shortages prevent females from reaching a required body condition threshold (Boyd et al. 1999). Such maturation delays have been reported for California sea lions (*Zalophus californianus*) during strong El Niño years (Melin et al. 2012), and Steller sea lions following the northeast Pacific marine heatwave (Maniscalco et al. 2026).

Energetic limitation may also elevate predation risk, particularly for mothers and calves (Srinivasan et al. 2018). When resources are limited, animals show increased acceptance of predation risk to maintain foraging intake (Dill and Gillett 1991). Under pre-heatwave conditions, Shark Bay dolphin foraging distribution reflected a trade-off between predation risk and food availability (Heithaus and Dill 2002), thus, reduced prey availability may force dolphins to forage in riskier habitats or for longer periods, increasing predation risk. Reduced vigilance and increased foraging effort in this context has previously been reported in Belding’s ground squirrels (*Spermophilus beldringi,* Bachman 1993), suggesting that the trade-off between energetic demand and predation risk may be a general response to resource limitation across taxa.

Reduced prey availability likely also had broader, population-level energetic consequences across Shark Bay, necessitating increased foraging effort while lowering energetic intake, in turn eroding body condition and, consequently, survival and reproductive output. Similar energetic trade-offs have been documented in Antarctic fur seals (*Arctocephalus gazella*), where increased foraging trip duration and energy expenditure during periods of low krill abundance resulted in lower maternal body mass and increased pup mortality (Costa et al. 1989). Nutritional stress impairs growth, reproduction (Williams et al. 2013; Wasser et al. 2017) and immune function (Raverty et al. 2020) in cetaceans, with consequences for individual fitness and population viability (Serres et al. 2024). Even modest fluctuations in prey availability can prevent marine mammals meeting energetic demands, with detrimental effects on body condition and survival (Stewart et al. 2021; Rojano-Doñate et al. 2024). Such reductions in energy intake have, in turn, been linked to population collapse in marine mammal species, such as harbour porpoises (*Phocoena phocoena*; Gallagher et al. 2022).

The sustained reduction in survival and recruitment drove prolonged population abundance declines in both gulfs. These losses were substantial: a 38.8% (36.1% - 41.2%) decline in WSB between 2011 and 2018 and a 31.6% (27.3% - 36.0%) decline in ESB between 2011 and 2023, comparable in magnitude to the 37% decline documented in Hawaiian humpback whales following the Northeast Pacific marine heatwave (Barber-Meyer et al. 2026). While WSB showed an immediate post-heatwave drop in abundance, ESB abundance remained stable until 2014, despite survival declining a year earlier. This pattern is consistent with the recognised decoupling between vital rate changes and population-level responses under density-independent disturbances (Bradshaw and Herrando-Pérez 2023). This underscores the risk of relying on population counts alone to assess vulnerability in long-lived species, as vital rate declines can precede detectable abundance changes by several years (Perrig et al. 2019).

The magnitude of the abundance reductions documented here is of particular concern. In other species with slow life histories and limited capacity for rapid population growth, such as North Atlantic right whales (*Eubalaena glacialisi*), post-decline recovery has been extremely slow (Fujiwara and Caswell 2001). Furthermore, recovery probability decreases the longer populations remain in a depleted state (Hutchings 2015), a dynamic relevant to ESB, where demographic decline has persisted for over a decade. While we did not assess genetic consequences directly, substantial reductions in population size can also erode genetic diversity and increase effects of genetic drift and inbreeding (Frankham 2005), which may reduce fitness (Reed and Frankham 2003), and reduce resilience to environmental change (Caughley 1994).

The observed spatial heterogeneity in impact and recovery trajectories between the gulfs was associated with differences in seagrass loss. Across study sites, 56% of pre-heatwave coverage (140 km^2^) was lost in WSB and 40% coverage (85 km^2^) in ESB. Seagrass decline was more rapid and extensive in WSB, and more gradual and sustained in ESB (Fig. 2; Supplementary Methods S2.3), a pattern that mirrors the demographic trajectories of each gulf. The smaller initial loss of seagrass in ESB may have buffered short-term impacts on prey availability, delaying the demographic response. Dolphins may target new species in response to declining abundance of preferred prey (Stolen et al. 2025), allowing short-term compensation for more gradual prey declines. However, the prolonged absence of recovery, and lower baseline seagrass coverage in ESB, may have driven prey declines past a critical threshold beyond which dolphins could no longer compensate.

Seagrass coverage, as the annual proportion of each study area, was positively associated with adult survival in WSB; the ESB estimate was also positive, but its 95% credible interval overlapped zero, with directionally consistent but weaker effects on juvenile and calf survival. Seagrass coverage in the year prior to conception also impacted reproductive output, showing a positive association with recruitment in ESB, although the 95% credible interval narrowly overlapped zero. The support for this pre-conception lag, rather than seagrass coverage during gestation or conception, suggests that maternal body condition entering a reproductive attempt was critical for successful recruitment. This is consistent with the energetic demands of dolphin reproduction, where gestation typically lasts 12 months, followed by an extended lactation period (Mann et al. 2000). In WSB, the relationship was directionally consistent but not significant, likely reflecting the shorter duration of severe habitat degradation. This pathway of demographic impacts through the loss of habitat forming species is likely to operate across diverse ecosystems, and become increasingly relevant as such losses become more frequent (Smale et al. 2019).

Vital rates remained depressed for several years following the heatwave in both gulfs. The strong, delayed effect of seagrass loss on recruitment points to habitat and prey effects that rippled through the ecosystem over extended periods. The extended timescale of these effects likely mirrors the slow recovery of *A. antarctica* (Hemminga and Duarte 2000), which can take decades (O’Brien et al. 2018) - a recovery further inhibited by ongoing post-heatwave warming observed (Marbà and Duarte 2010; Gilmour et al., 2019), particularly in ESB, where cMHW data show fewer returns to near-baseline conditions than WSB (Fig. 1B,C). While fast growing, typically tropical colonizing seagrasses increased post-heatwave (Kendrick et al. 2019), these are unlikely to replace the structural and functional roles of *A. antarctica* or compensate for the lost ecosystem services (Hyndes et al. 2003).

Despite concurrent depression of survival and recruitment, WSB showed some evidence of demographic resilience (Holling 1973; McKinney 1997), with positive growth rates and partial abundance recovery from 2019 onwards. In contrast, ESB showed no comparable recovery signal in abundance. Recovery trajectories in both gulfs tracked seagrass dynamics, suggesting resilience in this system may be primarily habitat-mediated, contingent on the pace and extent of ecosystem recovery rather than being an intrinsic property of the sub-populations. Such links between habitat recovery and recovery of dependent communities have been documented elsewhere. For example, the restoration of eelgrass meadows resulted in the rapid return of fish biomass and ecosystem services in inshore lagoons of Virginia, USA (Orth et al. 2020), while long-term studies of coral reef communities have shown that consumer assemblages closely track trajectories of habitat disturbance and recovery (Berumen and Pratchett 2006).

Such recovery can be constrained by the frequency of repeat disturbances, as populations may experience long-term decline when disturbance intervals are shorter than recovery times, even if resilient to individual events (Fairman et al. 2016; Neilson et al. 2020; Paine et al. 1998). For bottlenose dolphins (*T. truncatus*) inhabiting the Little Bahama Bank, repeated hurricane activity influenced apparent survival through increased mortality and/or permanent emigration (Coxon et al. 2022), with repeated hurricane exposure also proposed as a contributing factor to the continued decline of the neighbouring East Abaco population (Fearnbach et al. 2012). Our cumulative marine heatwave intensity data also indicates this dynamic, with sustained elevation in cMHW intensity post-2011 in ESB coinciding with continued decline. We defined these events using a fixed climatological baseline (1982-2011), which better captures ecologically relevant departures from historical conditions for species with slow adaptive responses to warming (Zhang et al. 2023). The sustained elevation in cMHW intensity in ESB reflects a combination of repeated discrete heatwave events and rising background temperatures relative to the baseline (Capotondi et al. 2024). Both represent a sustained departure from pre-heatwave conditions, suggesting that sub-catastrophic events, along with chronic thermal stress, may impede recovery in populations not yet returned to pre-disturbance vital rates. This is particularly concerning given the projected global increase in marine heatwave frequency, duration and intensity throughout the 21st century (Meehl and Tebaldi 2004).

Accurately characterising the demographic consequences of these events requires multi-decadal datasets spanning pre-disturbance baselines and post-disturbance trajectories. These datasets remain rare in marine research despite their necessity for detecting delayed and prolonged demographic responses and informing effective conservation management (Maxwell et al. 2019). Our 20-year dataset provided a uniquely well-characterised pre-heatwave baseline and captures the full demographic trajectory of decline and partial recovery. Critically, by monitoring two sub-populations exposed to the same disturbance but diverging in habitat recovery, we gain stronger evidence for a habitat-related explanation and a clear demonstration that aggregated, population-wide monitoring can mask acute conservation concerns playing out at finer spatial scales. This divergence points to foundation habitat protection as the most relevant management lever in this system.

We show that the 2011 marine heatwave inflicted demographic damage on dolphins that was severe and prolonged. More than a decade after the initiating event, the ESB sub-population remains in demographic decline, while recovery in WSB has only been partial. This is a stark result: extreme climate events can overwhelm the adaptive potential of even behaviourally flexible species, such as bottlenose dolphins. As marine heatwaves grow more frequent under projected climate change, the intervals between disturbances may become too short for long-lived megafauna with slow life histories to recover. Our study delivers rare, multi-decadal empirical evidence of the demographic costs of an extreme climate event and makes clear that conservation strategies must be built around habitat protection and climate assessments that account for the compounding toll of repeated disturbance.

## Supporting information

Supplementary Information for Smith et al 2026

## 5. Acknowledgements

This research was carried out in Gathaaguda, Malgana Country, and we respectfully acknowledge the Traditional Owners of the region. We thank RAC Monkey Mia Dolphin Resort, Shark Bay Resources, the Useless Loop community and the local Department of Biodiversity, Conservation and Attractions (DBCA) staff for their ongoing logistical support. We are extremely grateful to all members and field assistants of Shark Bay Dolphin Research (www.sharkbaydolphins.org) for their long-term contributions to data collection, without whom this research would not be possible. This study was funded by Swiss National Science Foundation grant (Grant Number 310030_204974) awarded to M.K., the A.H. Schultz Foundation, the University of Zürich. D.M.B. was funded by the Swiss National Science Foundation (Grant Number P500PB_225411).

## Notes

### Competing Interest Statement

The authors have declared no competing interest.

### Summary of Updates

Figures 4 and 7 have been revised

