## Supplementary Information for Smith et al 2026 for "Demographic change in an iconic dolphin population driven by habitat degradation following a marine heatwave"

**Supporting Information for Demographic change in an iconic dolphin population driven by habitat degradation following a marine heatwave**

**List of Authors:**

Felix Smith^+,^ Dominik M. Behr, Manuela R. Bizzozzero, Simon J. Allen, Richard C. Connor, Stephanie L. King, Arpat Ozgul, Michael Krützen

### Supplementary Methods

#### S1. Estimation of cumulative marine heatwave intensity

Cumulative marine heatwave intensity (°Cdays; Hobday et al., 2016) was calculated to represent an external driver of ecosystem change. While not directly included in the HMM model, it was used to infer seagrass coverage for years with missing data via an autoregressive sub-model described below (S2.2) Daily sea surface temperature (SST) data for Shark Bay were obtained from the NOAA Optimum Interpolation Sea Surface dataset (OISST; NOAA National Centers for Environmental Information) via the Environmental Research Division Data Access Program (ERDDAP) data server (dataset ID: ncdcOisst21Agg_LonPM180). The OISST product is a global ¼ degree gridded dataset of Advanced Very High Resolution Radiometer derived SSTs at daily resolution (Reynolds et al. 2007). SSTs were extracted for the period 01 January 1982 to 31 December 2023. The extracted OISST data comprised four spatial grid cells within each study site per day. For each site, we calculated the daily mean SST by averaging values across the four grid cells. Using the R package *heatwaveR* (Schlegel and Smit 2018), we constructed a 30-year historical climatology for each gulf of Shark Bay and identified marine heatwave events following the definition of Hobday et al., (2016). For each detected event, cumulative intensity was calculated. Annual cumulative intensity was derived as the sum of event-level cumulative intensities occurring during each biological year (01 June year *t* to 31 May year *t* + 1).

##### S1.2. Annual cumulative marine heatwave intensity

Cumulative marine heatwave intensity was low between 2002 and 2009 in both gulfs of Shark Bay (Figure S1). Both gulfs experienced an extreme peak in cMHW intensity in 2010, reaching 380°Cdays in WSB and 326.6°Cdays in ESB. In WSB, elevated intensity remained until 2012, after which intensities were moderate to low throughout the rest of the study period (14.8°Cdays-75.5°Cdays). In ESB, cMHW intensity remained high until 2015, before declining in 2016 (Figure S2). Secondary peaks were observed in 2019 (111.5°Cdays) and 2023 (160.6°Cdays).

A	B

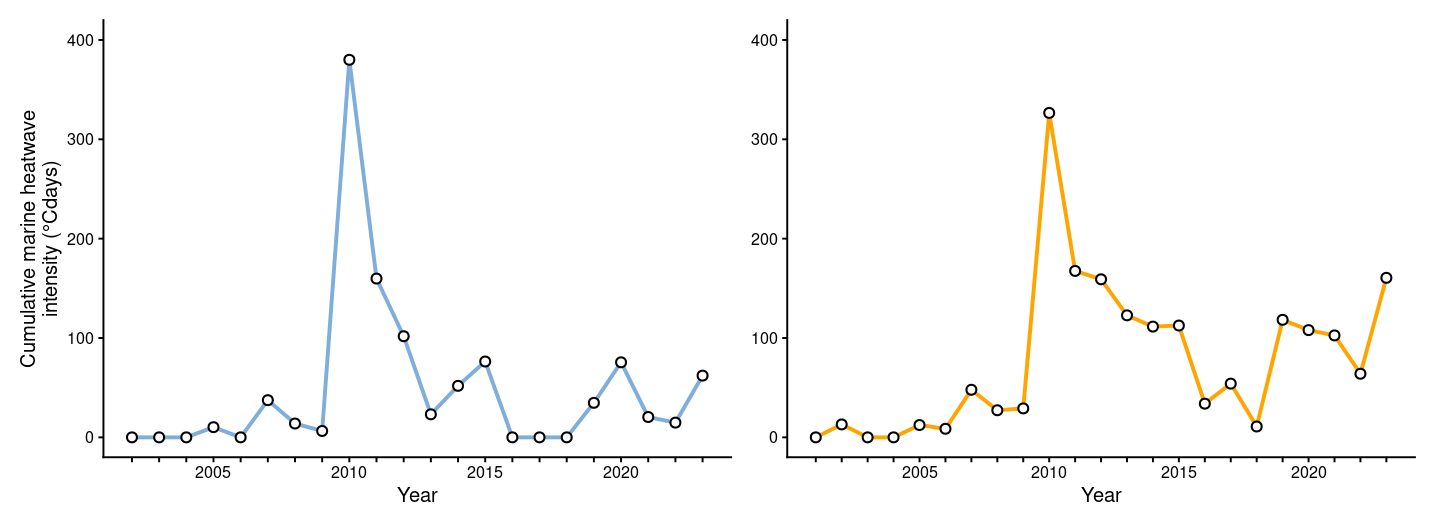

**Figure S1:** Annual cumulative marine heatwave intensity (°CDays) for West Shark Bay (A) and East Shark Bay (B). Cumulative marine heatwave intensity represents the sum of daily temperature anomalies (°C) above a fixed 30-year climatological threshold across all days classified as part of a marine heatwave event within each year

#### S2. Estimation of annual seagrass coverage

##### S2.1. Remote sensing and classification

We generated annual seagrass coverage maps following methods described in Bizzozzero et al. (2025). Annual seagrass layers were derived from Landsat imagery from 2001–2023, excluding 2011 and 2012 due to data availability constraints and sensor limitations reducing image quality during this period (Hossain et al. 2015; Nowicki et al. 2017). We used Landsat Surface Reflectant imagery (Landsat 5 Level 2, Collection 2, Tier 1; Landsat 7 Level 2, Collection 2, Tier 1; and Landsat 8, Collection 2, Tier 1; USGS, 2020) at 30 m resolution and including atmospheric correction. Imagery was selected from the period 1^st^ May to 30^th^ November each year, aligning with primary survey periods (see section 2.4.1 of main text). Annual mean high-quality composite images of RGB bands were generated in Google Earth Engine and classified into “seagrass” and “non-seagrass” pixels by applying a random forest algorithm using a custom JavaScript developed by Bizzozzero et al. (2025).

We performed a principal component analysis (PCA) on the composite imagery for the years 2001–2010 and 2013–2023 to reduce correlation of the input data layers used for classification. Each year, 233–283 points were classified as “seagrass” and 276–333 points as “other” based on visual inspection of the high-quality composite images. We trained a random forest algorithm on the PCA values of the classified pixels to identify pixels of indicative seagrass. Point locations were fixed across all years and were revalidated annually. Classification of points was validated using an independent dataset of 100 randomly selected points across the study area, which were not included in model training. These served as a consistent validation set across all years. We re-evaluated the same 100 points manually each year using the high-quality composite imagery, ensuring consistency in classification accuracy assessment across years. For each year, the total number of pixels classified as seagrass within each study site was converted to area (km²) based on the raster pixel resolution.

Classification performance of annual seagrass maps was evaluated using the Kappa coefficient (Landis and Koch 1977). Values above 0.61 are defined as “substantial agreement”, with those above 0.81 reflecting “almost perfect agreement”. Values ranging from 0.41-0.60 indicate “moderate agreement”. Kappa values ranged from 0.539 – 0.899, with an overall mean of 0.726, indicating substantial agreement between classified and reference data (Fig. S2).

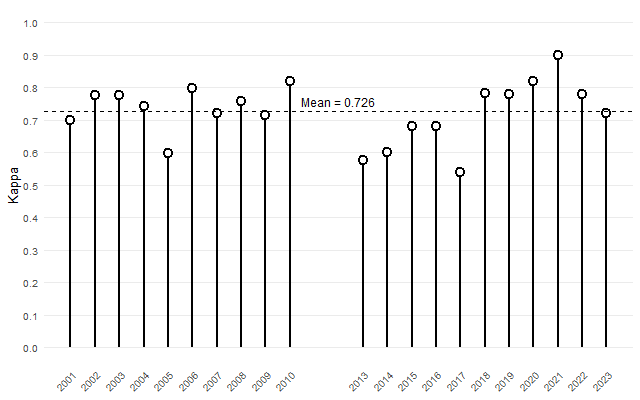

**Figure S2:** Validation Kappa values for annual seagrass classification maps from 2002 to 2019 (excluding 2011-2012). Points show Kappa coefficient for each year; dotted line represents the overall Kappa value (0.726). Higher values indicate greater agreement between classified and reference data.

##### S2.2. Seagrass sub-model

To impute seagrass coverage for missing years, we embedded a seagrass process sub-model within the Bayesian hidden Markov model. We treated missing seagrass values as latent variables and estimated these jointly with all other model parameters. The seagrass sub-model followed a first-order autoregressive AR(1) process informed by lagged cumulative marine heatwave intensity:

seagrass*_t_* ~ Normal(*α* + *β_cMHW_* log(cMHW*_t−1_*) + *ρ* seagrass*_t−1_*, *σ_sg_*)

where α is the intercept; *β_cMHW_* is the effect of log-transformed lagged cumulative marine heatwave intensity on seagrass coverage; *ρ* is the first-order autoregressive coefficient capturing temporal persistence in seagrass coverage; and σ_sg_ is the residual standard deviation.

We log transformed cumulative marine heatwave intensity to reduce the influence of extreme values (e.g., the 2011 marine heatwave event) on predictions. For years with observed seagrass data, the data informed the sub-model likelihood, effectively constraining these values to their observed levels. For the two missing years, the sub-model generated posterior predictions informed by the temporal autocorrelation structure and the known heatwave conditions during that period.

##### S2.3. Temporal variation in seagrass coverage

Seagrass extent within the WSB study area remained relatively stable prior to the 2011 marine heatwave (Figure S3) with an average coverage of 247.4 km^2^ between 2006 and 2010 (37.6% cover). Seagrass coverage declined sharply between 2010 and 2011, reaching a model-derived coverage of 111.3 km^2^ (16.9%) and continuing to decline in 2012 (107.3 km^2^; 16.3%). Coverage remained low until 2016 (117.8 km^2^; 17.8%). A marked recovery occurred between 2017 and 2018, before a subsequent decline to 118.2km^2^ (17.9%) in 2023.

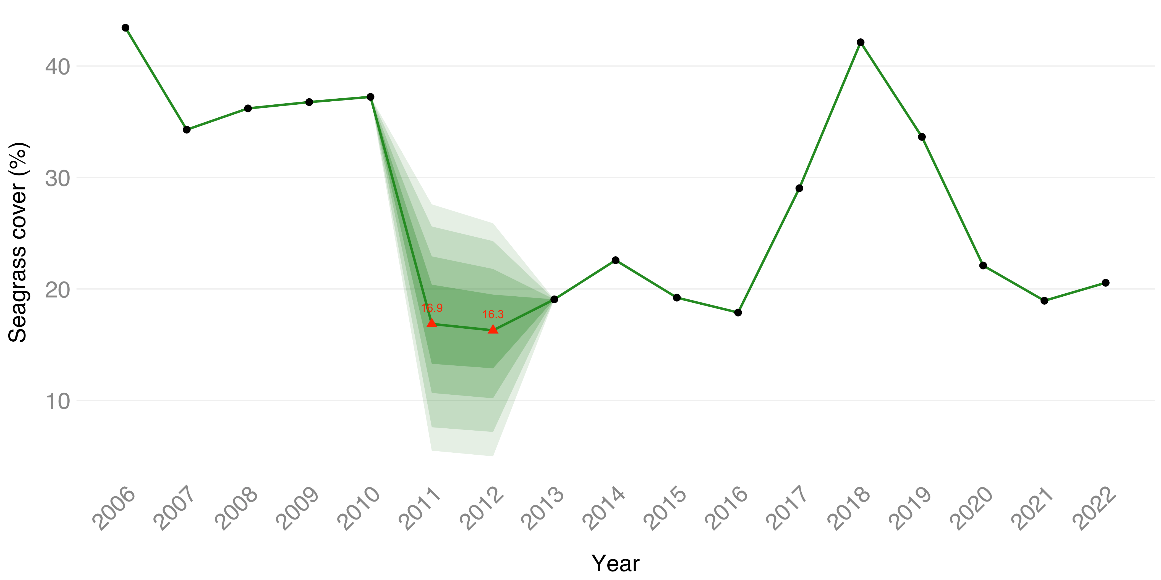

**Figure S3:** Annual seagrass cover (%) in West Shark Bay. Black dots = observed data, red triangles = imputed values from the seagrass sub-model. Ribbons show graduated posterior credible intervals (50%, 75%, 90%, 95%).

Seagrass coverage overall was lower in ESB than WSB. Within the ESB study area, seagrass extent remained generally consistent between 2002 and 2010 around an average coverage of 178.1 km^2^ (19.8% coverage; Figure S4), with some interannual variability including a peak in 2006. Following the 2011 marine heatwave there was a steady decline in seagrass coverage reaching a minimum of 106.1 km^2^ in 2019 (11.8% coverage). This was followed by modest recovery and a subsequent decline to 120.3 km^2^ in 2023 (13.4% coverage).

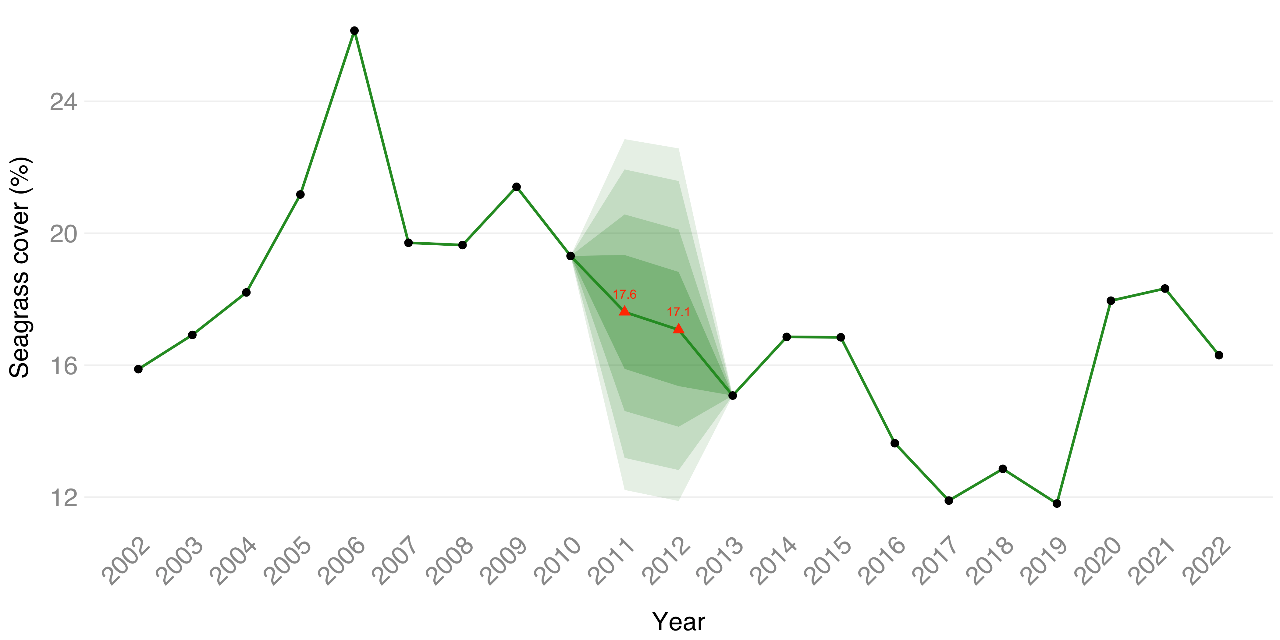

**Figure S4:** Annual seagrass cover (%) in Eest Shark Bay. Black dots = observed data, red triangles = imputed values from the seagrass sub-model. Ribbons show graduated posterior credible intervals (50%, 75%, 90%, 95%).

#### S3. Consideration of mark recapture model assumptions

Several model assumptions and study-specific factors should be considered (Pollock et al. 1990; Williams et al. 2002). Firstly, marks should be unique, permanent and not mis-identified. The acquisition of marks is cumulative in dolphins (Wursig and Jefferson 1990), meaning younger animals are often less distinctively marked than adults. Failure to recognise previously marked dolphins may bias capture and survival rates downward, with an upward bias in abundance (Silva et al. 2009). To avoid mis-identification, we used supplementary identification features, such as fin shape and scarring patterns, when individuals were less distinctive and all photo-identification data were double-checked by a second observer.

Secondly, all individuals have equal probability of being captured in a given secondary period. This is rarely true for cetacean populations (Hammond 1986), as individual detection varies with age, sex, social structure and spatial use (Nicholson et al. 2012; Pollock et al. 1990). We integrated age-class-specific detection probabilities and a spatial effort covariate to control for age-class and spatial heterogeneity in detection.

Individual capture probability should also be independent of others. The social structure of dolphins in Shark Bay (Connor et al. 2000) mean individuals are frequently detected in association with specific conspecifics, potentially violating this assumption. Abundance estimates are relatively robust to such violations however, although model-based variances may be underestimated (Williams et al. 2002).

Finally, the population should remain closed during primary periods. Demographic closure is unlikely to be strictly met in natural populations (e.g.., Silva et al., 2009), although several features of our study reduce the likelihood of substantial bias. Primary periods were short relative to the temporal scale of demographic processes, movement between the study sites is known to be negligible (Krützen et al. 2004) and previous work suggests no evidence of closure violations in this population (Rankin et al. 2016).

#### S4. Parameterization of the Hidden Markov Model

Individuals occupy latent biological states that represent the individual’s true state during primary period *t*, defined by age-class (*calf, juvenile, adult*) and location (*onsite, offsite*), with additional states for *not yet entered the population* and *dead*. To parameterise the HMM-robust design model, we employed a multistate Jolly-Seber formulation incorporating both age-class and individual location. Transitions between latent states from *t* to *t+1* were governed by a state-transition matrix **Ω** (Table S1) comprising the demographic parameters survival (Φ), temporary emigration (γ′ and γ″), and entry processes (ω and λ). To account for imperfect detection, we modelled the observation process alongside the state process (Kéry and Schaub 2012). Observed states were linked to true latent states through a probabilistic observation matrix **Θ** (Table S2) that specified the likelihood of detecting an individual in a given observation category, conditional on its true state in year *t*. For each individual *i*, the sequence of latent states *z_i,t_* was modeled hierarchically based on the state transition matrix **Ω**, the observation matrix **Θ**, and the detection history *Y_i,t,s_* as

$z_{i,t+1} | z_{i,t} \sim Categorical\left( \boldsymbol{\Omega}_{z_{i,t},t,1\cdots S} \right)$ and

$Y_{i,t,s} | z_{i,t} \sim Categorical\left( \boldsymbol{\Theta}_{z_{i,t},i,t,s,1\cdots O} \right)$,

where *S* denotes the number of latent states, and *O* the number of observed states.

We modeled full capture histories (rather than conditioning on first capture) to allow inference about state transitions and sub-population rate of change (Rankin et al. 2016). To estimate the proportion of recruits entering onsite versus offsite, we used the eigenvector decomposition method of Rankin et al. (2016). This approach assumes that recruits enter the onsite and offsite states in proportions consistent with the temporary emigration process of the marked population, an assumption we consider reasonable for Shark Bay dolphins, as suitable habitat

**Table S1:** State-transition matrix **Ω** for bottlenose dolphins in Shark Bay. *ω_t_* denotes the year-dependent entry probability, defined as the probability that an individual not yet entered into the population transitions into the population in year *t*. *λ* denotes the proportion of newly entered individuals that start onsite and are available for detection. ${\Phi X}_{i,t}$ is the age-specific survival of individual *i* in primary period *t.* Age-class transitions are governed by $a_{i,t}^{X}$ which equals 1 if an individual *i* at time *t* advances to the next age class at time *t* + 1, and 0 otherwise. γ' represents the probability of moving offsite at time *t* + 1 given the individual was onsite at time *t*; γ'' represents the probability of staying offsite at time *t* + 1 given the individual was offsite at time *t.*

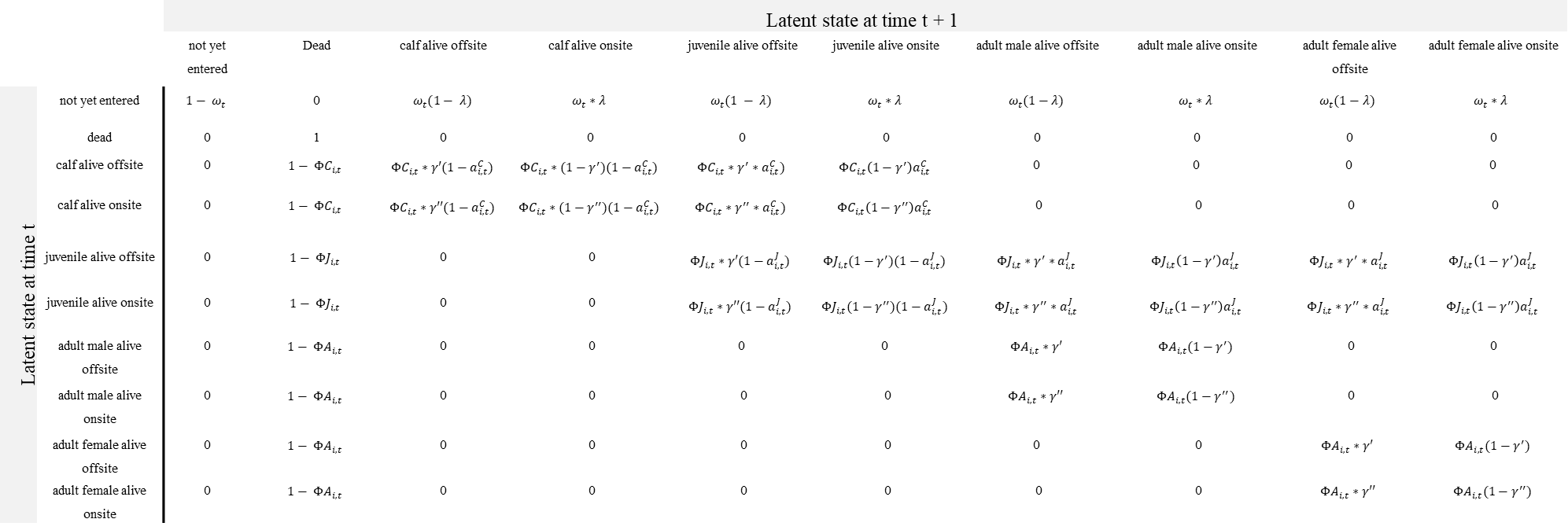

extends continuously beyond the boundaries of each study area and individuals can move freely between the surveyed and unsurveyed portions of the gulf (Preen et al. 1997).

To account for uncertainty related to age of individual dolphins, we estimated birthdate as a latent variable for each animal from an individual-specific normal distribution centred on its estimated birthdate with a standard deviation derived from the birthdate uncertainty divided by 1.96. Age-class was then determined based on the sampled birth date. Similarly, the sex for unsexed dolphins was inferred as a latent variable via a Bernoulli draw parameterized by the empirically observed proportion of females in the respective gulf (Bond et al. 2025).

**Table S2:** Observation matrix **Θ** for bottlenose dolphins in Shark Bay. *pX_i,t,s_* denotes the individual- (*i*), year- (*t*), and secondary period- (*s*) dependent probability of detection for calves (*X* = *C*), juveniles (*X* = *J*), and adults (*X* = *A*); *δ_t_* indicates whether surveys were conducted in year *t*; *π_i,t,s_* is individual *i*'s availability (zone-based) during secondary *s*.

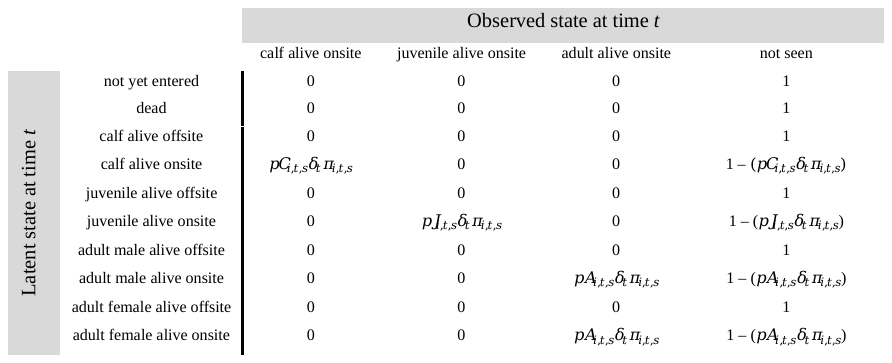

##### S4.1. Modelling detection probability

We modelled detection probability to account for imperfect and heterogenous detection driven by variation in survey effort across individuals, space, sampling periods and age-class. The detection probability of individual *i* during secondary period *s* within primary period *t* was modelled by Bernoulli trial with age-class specific success probability *p_i,t,s_.* Detection was conditional on the individual being alive and available for detection in any of the study area zones surveyed during that secondary period (see S5). We modelled individual-, year- and secondary occasion-dependent detection probabilities as:

$$Logit\left( pC_{i,t,s} \right)= a_{1}+ \alpha_{4}{effort}_{i,t,s}+ {\varepsilon C}_{t}$$

$$Logit\left( pJ_{i,t,s} \right)= a_{2}+ \alpha_{4}{effort}_{i,t,s}+ {\varepsilon J}_{t}$$

$$Logit\left( pA_{i,t,s} \right)= a_{3}+ \alpha_{4}{effort}_{i,t,s}+ {\varepsilon A}_{t}$$

Here C = calves, J = juveniles, and A = adults; $a_{1}, a_{2}$ and $a_{3}$ are the age-class-specific intercepts of detection; $a_{4}$ is the linear effect of survey effort on logit of detection (irrespective of age-class); ${effort}_{i,t,s}$ is the individual-specific survey effort undertaken in the survey zone occupied by individual *i* during secondary period *s* of primary period *t,* calculated from the proportion of days the survey zone the individual occupied was surveyed during that secondary period; and $\varepsilon_{t}$ is the age-class-specific random effect of year on the intercept of detection, with $\varepsilon_{t}$ ~ Normal(0, σ). Detection probabilities were fixed to 0 in years with no survey effort (2007 and 2008 in ESB), indicated by δ*_t_* (δ*_t_* =0 for years with no survey effort, and δ*_t_* =1 otherwise) in the observation matrix (Table S2). Individual availability for detection was conditioned on zone occupancy, with unobserved individuals assigned to zones based on their historical detection frequency. Full details are provided in Supplementary Methods S5.

##### S4.2. Modelling survival and temporary emigration probabilities

We modelled survival and temporary emigration probabilities to quantify demographic responses to the marine heatwave, while distinguishing true mortality from temporary absence due to movement outside of the study area. Survival probability represented the probability of an individual alive in year *t* surviving to year *t* + 1. Survival of an individual *i* started upon entry into the population and was subsequently modelled through Bernoulli trials with age-class specific success probability *ΦX_i,t_*(*t*=1,…,*T*-1). Individual- and year-specific survival probabilities were modelled as:

$$Logit\left( {\Phi C}_{i,t} \right)= \beta_{1}+ \beta_{2} \mathrm{seagrass}_{t}+{\varepsilon C}_{t}$$

$$Logit\left( {\Phi J}_{i,t} \right)= \beta_{3}+ \beta_{4}\mathrm{seagrass}_{t}+{\varepsilon J}_{t}$$

$$Logit\left( {\Phi A}_{i,t} \right)= \beta_{5}+ \beta_{6}\mathrm{seagrass}_{t}+\beta_{7}{forage}_{i}+ \beta_{8}{sex}_{i}+{\varepsilon A}_{t}$$

Here, $\beta_{1}$, $\beta_{3}$, and $\beta_{5}$ are the age-specific intercepts corresponding to the mean survival probability on the logit scale; $\beta_{2}, \beta_{4}$ and $\beta_{6}$ are the age-class-specific linear effects of seagrass coverage of the current year on survival; $\beta_{7}$ is the linear effect of forage type (1 = sponger, 0 = non-sponger) on the logit of adult survival; $\beta_{8}$ is the linear effect of sex (1=female, 0=male) on the logit of adult survival (the effects of forage type and sex on survival were not modelled for younger age classes); and ${\varepsilon X}_{t}$ is the age-class-specific random effect of year on survival, with ${\varepsilon X}_{t}$ ~ Normal(0, σ). In all analyses, continuous covariates were standardised by subtracting their mean and dividing by their standard deviation.

We originally evaluated two alternative environmental covariate structures for survival: seagrass coverage at time *t* (current year) and seagrass coverage at time *t−1* (one-year lag). The former model provided the better model fit for both populations and was retained as the final environmental covariate structure. Due to strong temporal autocorrelation of seagrass coverage in both gulfs, we did not include a two-way interaction between seagrass coverage of the current and the previous year.

Using the Markovian formulation of Pollock's robust design (Kendall et al. 1997), we modelled temporary emigration with two time-constant parameters shared across all age classes: γ′, the probability of moving offsite given onsite, and γ″, the probability of remaining offsite given offsite in the preceding primary period. We assigned uniform priors constrained to γ′ ~ U(0, 0.5) and γ″ ~ U(0.5, 1), reflecting the biological assumption that temporary absence is a persistent state (animals offsite are more likely to remain offsite than to return within a single primary period) and ensuring that the onsite and offsite states remain distinguishable.

##### S4.3. Estimation of derived population-level demographic parameters

In addition to survival and detection probabilities, we derived age-class specific annual abundance, recruitment and population growth rate to quantify population-level responses to the marine heatwave. Abundance (*N_t_)* was defined as the total number of individuals alive at time *t* and included both onsite and offsite dolphins. We used data augmentation to account for animals that likely entered the subpopulation but were never detected (Royle and Dorazio 2012). To implement this, we added a set of pseudo-individuals with all-zero capture histories to the dataset. This approach accounts for imperfect detection that would otherwise lead to an underestimation if only observed capture histories were used. For each gulf the number of pseudo-individuals was equal to the total number of observed dolphins (ESB: *n* = 1197, WSB: *n* = 780). The model used estimated detection probabilities to infer which of these individuals represent real but unobserved animals. The true population size is therefore the sum of the observed individuals and those inferred to be real among the augmented pool. We assigned sex to pseudo-individuals via Bernoulli trial based on the empirically observed proportion of females in each sub-population, as per the unsexed observed individuals (Bond et al. 2025). Similarly, pseudo-individuals were assigned age based on birthdate estimates and uncertainties from observed individuals. This ensured pseudo-individuals reflected the sex ratio, age structure and associated uncertainty of the observed population. Annual population growth rate was calculated as the total number of animals alive at time *t*, divided by the total number alive at time *t*-1.

Recruitment rate represented the number of calves entering the population in year *t* relative to the number of adult females alive at year *t –* 1, where calves have survived long enough to be observed in the study. To estimate annual recruitment rate we used posterior estimates of the number of adult females and annual births from the Bayesian HMM, with full propagation of posterior uncertainty in both quantities. We modelled the number of births in year *t* as a binomial outcome with the number of adult females in year *t* – 1 representing the number of trials, as

$b_{t} \sim Binomial(F_{t-1},r_{t})$,

where $b_{t}$ is the number of births in year *t*, $F_{t-1}$ is the number of adult females alive in year *t* – 1; and $r_{t}$ is the recruitment rate in year *t.* Recruitment rate was then modelled on the logit scale as function of seagrass coverage and a random year effect:

$logit\left( r_{t} \right)= \alpha+ \beta_{k}\text{ seagrass}_{t-k}+ \varepsilon_{t}$,

where $\alpha$ is the intercept representing baseline recruitment rate on the logit scale, and $\varepsilon_{t}$ is a random year effect capturing unexplained annual variation, with $\varepsilon_{t}$ ~ Normal(0, σ). Due to high collinearity arising from temporal autocorrelation in seagrass coverage, we tested three biologically motivated time lags (*k* = 0, 1, 2) in separate models. Seagrass coverage in the survey year (*k* = 0) reflects food availability during gestation and at the time calves are first observed. The one-year lag (*k* = 1) reflects conditions during the conception year, which may directly affect pregnancy success. The two-year lag (*k* = 2) reflects pre-conception conditions and captures potential indirect effects on reproductive success via maternal body condition.

#### S5. Survey effort and zone occupancy

To account for unequal spatial survey effort within each study area across years, we subdivided each survey area (ESB and WSB) into survey zones (seven in ESB, eight in WSB) based on a combination of habitat zones described in Bizzozzero et al. (2025) and focal areas that were typically targeted on a given survey day (Fig. S5). For each survey zone within each secondary sampling period, we recorded the number of days that zone was surveyed, and used the proportion of days surveyed out of a possible 14 as a continuous effort covariate on detection probability (e.g., a zone surveyed on 7 of 14 days contributes an effort value of 0.5). Finally, for each individual detected within a specific secondary period, we recorded the survey zone in which the individual was observed, allowing us to account for individual-specific spatial heterogeneity in detection probability (see Section 2.4.2 of main text).

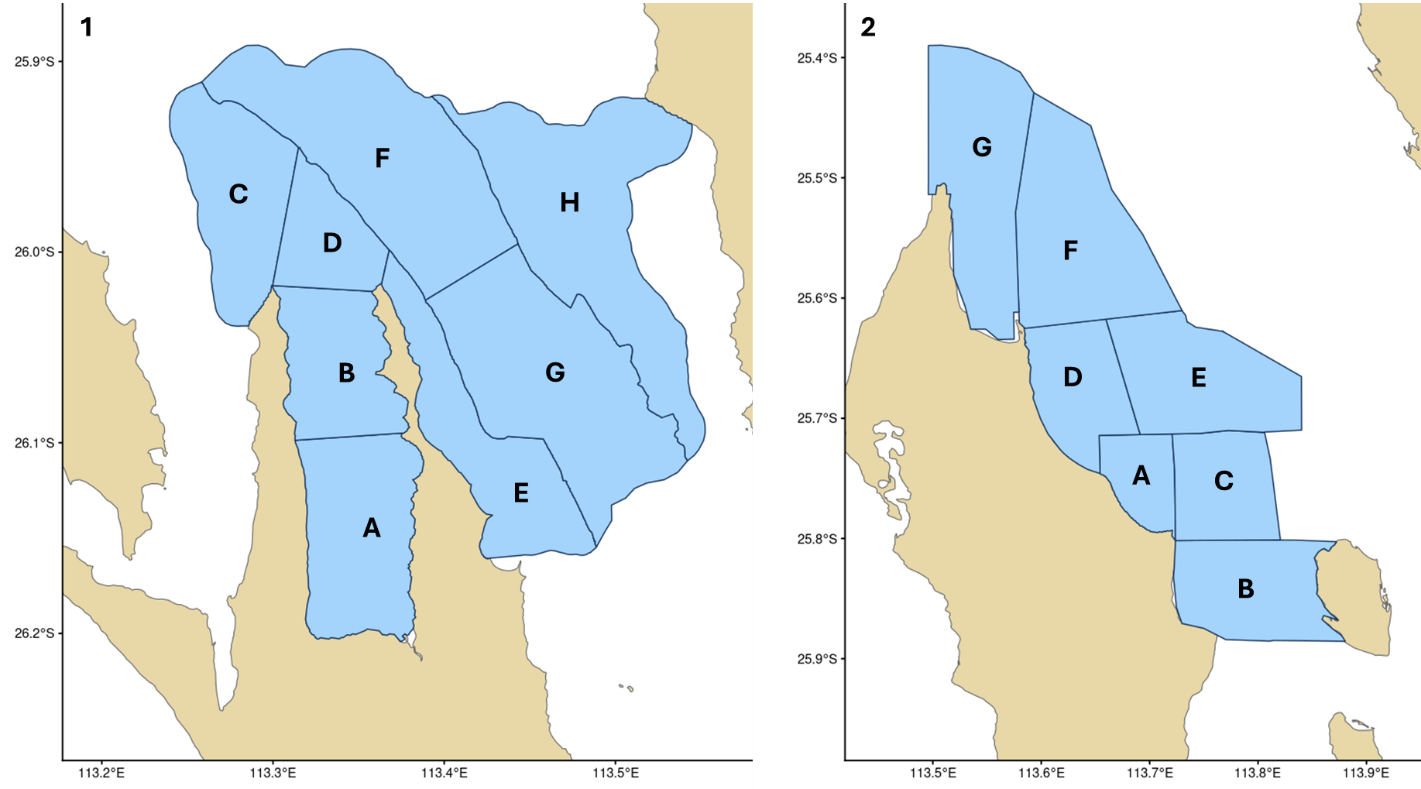

**Figure S5.** Survey Zones in West Shark Bay (1) and East Shark Bay (2). Zones represent focal areas typically targeted on a given survey day.

To account for individual movement between survey zones within a primary period, we conditioned an individual *i*’s availability for detection during a secondary period *s* (denoted by π_i,t,s_ in the observation matrix, see Supplementary Methods S4) on its zone occupancy during that period. When an individual was observed during a secondary period, its zone was known directly. When unobserved, zone occupancy was treated as a latent variable drawn from a categorical distribution reflecting that individual’s relative frequency of historical occurrence across zones. An individual was then considered available for detection only during secondary periods in which its occupied zones were surveyed. For example, if an individual was recorded in survey zone B during 80% of its detections, it would be available for detection, with probability 0.8 during secondary periods in which the individual was not observed but zone B was surveyed.

### Supplementary Results

#### S6. Assessment of Model Fit

We assessed model fit via posterior predictive checks. Across primary survey events, average Bayesian p-values of 0.77 in WSB and 0.65 in ESB indicated good overall fit. (Fig. S6). Anomalies observed in 2020 in WSB, reflecting near-zero survey coverage due to COVID-19 lockdowns, and 2007-2008 in ESB, where no surveys took place, are not indicative of model misfit. Similarly, the consistent over-prediction of detections in the earliest survey years of both gulfs is an expected feature of open-population models: individuals that were first identified later in the study (e.g., as adults or juveniles) are inferred to have been present earlier, inflating the modelled population available for detection before the photo-identification catalogue was saturated and while early survey coverage was sparse. Because our model allows only newborns to enter the population after the first year, such individuals are assumed to have been present from the outset, which would bias early abundance upward only if they had instead immigrated as adults. However, immigration of adults into either gulf is unlikely. Genetic analyses show that Shark Bay dolphins are markedly differentiated from populations to the north and south (Marfurt et al. 2024, 2026), consistent with limited historical gene flow and restricted dispersal into the study area. Early-year abundance estimates are therefore unlikely to be biased in either gulf.

To assess whether the elevated average Bayesian p-values reflected these boundary years, rather than systematic model misfit, we recomputed the mean over a restricted subsample of survey years. Exclusions were defined *a priori* on structural and effort ground, independently of model fit: we removed the first four survey years (the period during which open-population models inherently over-predict detections) and years with no or near-zero survey effort (2007-2008 in ESB; 2020 in WSB). The restricted mean Bayesian p-value moved toward the expected value of 0.5 in both gulfs, closely in ESB (0.51) and more modestly in WSB (0.69).

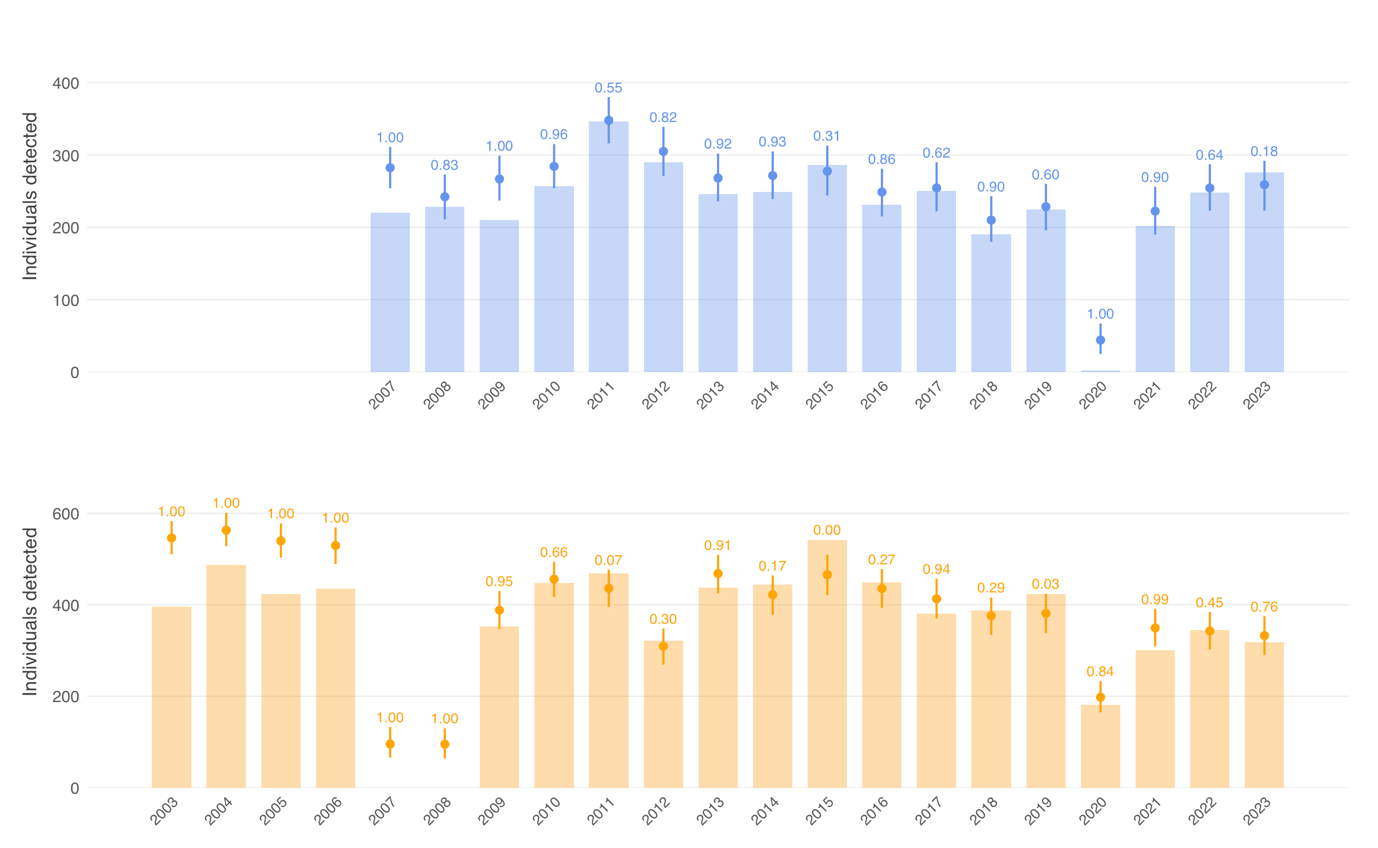

**Figure S6**: Posterior predictive check of the Bayesian HMM for the Western Shark Bay dolphin population (top panel) and the East Shark Bay population (bottom panel). Bars represent total number of observed individuals in each year, while points and error bars show the mean and 95% CIs of 1,000 replicated datasets simulated under the Bayesian HMM. The numbers above the error bars represent Bayesian p-values for the corresponding years, with values close to 0.5 indicating a good model fit (Gelman et al. 2013) and values outside the 0.10-0.90 range indicating poor fit (Hobbs and Hooten 2015).

#### S7. Detection probability estimates

Detection probabilities were similar across age classes in both WSB and ESB. While some moderate interannual variation in detection probability was observed, there was no evidence of extreme temporal variation. Detection probability increased strongly with survey effort (WSB: *ß* = 3.16, sd = 0.12; ESB: *ß* = 3.13, sd = 0.06), indicating variation in detectability was well captured by the effort covariate (Table S3).

**Table S3:** Mean, standard deviation and credible intervals of age-class-specific (calf < 3 yr, juvenile 3-12 yr, adult >12 yr) alpha parameters and random year effects for probability of detection in West and East Shark Bay. Ȓ of < 1.1 indicates convergence of MCMC chains. N.eff is effective sample size. All parameter estimates are on the inverse logit scale.

|  | **Parameter** | **Mean** | **SD** | **2.5%** | **50%** | **97.5%** | ***Ȓ*** | **n.eff** |
| --- | --- | --- | --- | --- | --- | --- | --- | --- |
| **West Shark Bay** | Calf detection probability | -1.351 | 0.069 | -1.486 | -1.352 | -1.213 | 1.05 | 5965 |
|  | Juvenile detection probability | -1.141 | 0.102 | -1.346 | -1.140 | -0.943 | 1.01 | 3825 |
|  | Adult detection probability | -1.302 | 0.098 | -1.499 | -1.303 | -1.104 | 1.04 | 1184 |
|  | Random year effect Calf | 0.164 | 0.066 | 0.057 | 0.157 | 0.313 | 1.00 | 2608 |
|  | Random year effect Juvenile | 0.339 | 0.088 | 0.203 | 0.327 | 0.544 | 1.01 | 9954 |
|  | Random year effect Adult | 0.350 | 0.084 | 0.227 | 0.337 | 0.551 | 1.01 | 9400 |
|  | Survey effort | 3.178 | 0.119 | 2.940 | 3.179 | 3.406 | 1.18 | 3297 |
| **East Shark Bay** | Calf detection probability | -1.623 | 0.087 | -1.791 | -1.624 | -1.448 | 1.00 | 3752 |
|  | Juvenile detection probability | -1.538 | 0.070 | -1.673 | -1.536 | -1.399 | 1.00 | 2814 |
|  | Adult detection probability | -1.538 | 0.057 | -1.649 | -1.538 | -1.426 | 1.01 | 1383 |
|  | Random year effect Calf | 0.324 | 0.077 | 0.201 | 0.314 | 0.500 | 1.01 | 7517 |
|  | Random year effect Juvenile | 0.269 | 0.056 | 0.181 | 0.262 | 0.398 | 1.00 | 12105 |
|  | Random year effect Adult | 0.221 | 0.042 | 0.154 | 0.215 | 0.320 | 1.00 | 14195 |
|  | Survey effort | 3.134 | 0.059 | 3.018 | 3.314 | 3.249 | 1.06 | 14450 |

#### S8. Temporary Emigration Parameters

Temporary emigration followed a Markovian process in both gulfs, with strong state persistence indicating consistent patterns of site use (Table S4). In both gulfs the probability of leaving the study area was low, reflecting strong site fidelity. However, individuals that were absent showed a high probability of remaining so in subsequent primary periods, suggesting that temporary emigration, when it occurred, tended to be persistent rather than transient.

**Table S4:** Temporary emigration parameters for WSB and ESB. γ' and γ'' parameterize the Markovian temporary emigration process. λ represents the probability of being onsite at time 1 and is derived from the equilibrium of the γ'/γ'' transition matrix: λ = (1 - γ'')/(1- γ' + 1- γ'')

|  |  | **Description** | **Mean** | **SD** | **2.5%** | **50%** | **97.5%** | **Rhat** | **n.eff** |
| --- | --- | --- | --- | --- | --- | --- | --- | --- | --- |
| **West Shark Bay** | $\gamma'$ | Probability of remaining offsite given offsite at time *t*-1 | 0.705 | 0.013 | 0.679 | 0.706 | 0.731 | 1.01 | 3953 |
|  | $\gamma''$ | Probability of moving offsite given onsite at time *t*-1 | 0.129 | 0.007 | 0.115 | 0.129 | 0.143 | 1.01 | 20234 |
|  | $\lambda$ | Probability of being available for capture in first primary period (initial site fidelity) | 0.696 | 0.012 | 0.673 | 0.696 | 0.719 | 1.01 | 4931 |
| **East Shark Bay** | $\gamma'$ | Probability of remaining offsite given offsite at time *t*-1 | 0.742 | 0.010 | 0.722 | 0.742 | 0.761 | 1.00 | 5172 |
|  | $\gamma''$ | Probability of moving offsite given onsite at time *t*-1 | 0.124 | 0.005 | 0.114 | 0.124 | 0.134 | 1.01 | 16070 |
|  | $\lambda$ | Probability of being available for capture in first primary period (initial site fidelity) | 0.675 | 0.009 | 0.657 | 0.675 | 0.693 | 1.00 | 5123 |

Kéry, Marc, and Michael Schaub. 2012. *Bayesian Population Analysis Using WinBUGS: A Hierarchical Perspective*. 1st ed. Academic Press.

Krützen, Michael, William B. Sherwin, Per Berggren, and Nick Gales. 2004. ‘POPULATION STRUCTURE IN AN INSHORE CETACEAN REVEALED BY MICROSATELLITE AND mtDNA ANALYSIS: BOTTLENOSE DOLPHINS’. *Marine Mammal Science* 20 (January): 28–47.

Landis, J. Richard, and Gary G. Koch. 1977. ‘An Application of Hierarchical Kappa-Type Statistics in the Assessment of Majority Agreement among Multiple Observers’. *Biometrics* 33 (2): 363. https://doi.org/10.2307/2529786.

Marfurt, Svenja M., Delphine B. H. Chabanne, Samuel Wittwer, et al. 2024. ‘Demographic History and Adaptive Evolution of Indo‐Pacific Bottlenose Dolphins ( *Tursiops Aduncus* ) in Western Australia’. *Molecular Ecology* 33 (22): e17555. https://doi.org/10.1111/mec.17555.

Marfurt, Svenja M., Adrien Tran Lu Y, Benjamin Dauphin, et al. 2026. ‘Whole‐Genome‐Sequencing Reveals Demographic History and Patterns of Parallel Adaptive Evolution in Indo‐Pacific Bottlenose Dolphins ( *Tursiops Aduncus* ) Across Coastal Australian Seascapes’. *Molecular Ecology* 35 (11): e70383. https://doi.org/10.1111/mec.70383.

Nicholson, Krista, Lars Bejder, Simon J. Allen, Michael Krtzen, and Kenneth H. Pollock. 2012. ‘Abundance, Survival and Temporary Emigration of Bottlenose Dolphins (Tursiops Sp.) off Useless Loop in the Western Gulf of Shark Bay, Western Australia’. *Marine and Freshwater Research* 63 (11): 1059–68. https://doi.org/10.1071/MF12210.

Nowicki, Rj, Ja Thomson, Da Burkholder, Jw Fourqurean, and Mr Heithaus. 2017. ‘Predicting Seagrass Recovery Times and Their Implications Following an Extreme Climate Event’. *Marine Ecology Progress Series* 567 (March): 79–93. https://doi.org/10.3354/meps12029.

Pollock, Kenneth H., James D. Nichols, Cavell Brownie, and James E. Hines. 1990. ‘Statistical Inference for Capture-Recapture Experiments’. *Wildlife Monographs* 107: 3–97.

Preen, A. R., H. Marsh, I. R. Lawler, R. I. T. Prince, and R. Shepherd. 1997. ‘Distribution and Abundance of Dugongs, Turtles, Dolphins and Other Megafauna in Shark Bay, Ningaloo Reef and Exmouth Gulf, Western Australia’. *Wildlife Research* 24 (2): 185–208. https://doi.org/10.1071/WR95078.

Rankin, Robert W., Krista E. Nicholson, Simon J. Allen, Michael Krützen, Lars Bejder, and Kenneth H. Pollock. 2016. ‘A Full-Capture Hierarchical Bayesian Model of Pollock’s Closed Robust Design and Application to Dolphins’. *Frontiers in Marine Science* 3 (March). https://doi.org/10.3389/fmars.2016.00025.

Reynolds, Richard W., Thomas M. Smith, Chunying Liu, Dudley B. Chelton, Kenneth S. Casey, and Michael G. Schlax. 2007. ‘Daily High-Resolution-Blended Analyses for Sea Surface Temperature’. *Journal of Climate* 20 (22): 5473–96. https://doi.org/10.1175/2007JCLI1824.1.

Royle, J. Andrew, and Robert M. Dorazio. 2012. ‘Parameter-Expanded Data Augmentation for Bayesian Analysis of Capture–Recapture Models’. *Journal of Ornithology* 152 (S2): 521–37. https://doi.org/10.1007/s10336-010-0619-4.

Schlegel, Robert W., and Albertus J. Smit. 2018. ‘heatwaveR: A Central Algorithm for the Detection of Heatwaves and Cold-Spells’. *Journal of Open Source Software* 3 (27): 821. https://doi.org/10.21105/joss.00821.

Silva, Ma, S. Magalhães, R. Prieto, Rs Santos, and Ps Hammond. 2009. ‘Estimating Survival and Abundance in a Bottlenose Dolphin Population Taking into Account Transience and Temporary Emigration’. *Marine Ecology Progress Series* 392 (October): 263–76. https://doi.org/10.3354/meps08233.

Williams, Byron K., James D. Nichols, and Michael James Conroy. 2002. *Analysis and Management of Animal Populations*. Academic press.

Wursig, B., and T. Jefferson. 1990. ‘Methods of Photo-Identification for Small Cetaceans’. *Reports of the International Whaling Commission*.
